# An Organism-Level Quantitative Flux Model of Energy Metabolism in Mice

**DOI:** 10.1101/2024.02.11.579776

**Authors:** Bo Yuan, Will Doxsey, Özlem Tok, Young Yon Kwon, Karen E. Inouye, Gökhan S. Hotamışlıgil, Sheng Hui

## Abstract

Mammalian tissues feed on nutrients in the blood circulation. At the organism-level, mammalian energy metabolism comprises of oxidation, storage, interconverting, and releasing of circulating nutrients. Though much is known about the individual processes and nutrients, a holistic and quantitative model describing these processes for all major circulating nutrients is lacking. Here, by integrating isotope tracer infusion, mass spectrometry, and isotope gas analyzer measurement, we developed a framework to systematically quantify fluxes through these metabolic processes for 10 major circulating energy nutrients in mice, resulting in an organism-level quantitative flux model of energy metabolism. This model revealed in wildtype mice that circulating nutrients have more dominant metabolic cycling fluxes than their oxidation fluxes, with distinct partition between cycling and oxidation flux for individual circulating nutrients. Applications of this framework in obese mouse models showed on a per animal basis extensive elevation of metabolic cycling fluxes in ob/ob mice, but not in diet-induced obese mice. Thus, our framework describes quantitatively the functioning of energy metabolism at the organism-level, valuable for revealing new features of energy metabolism in physiological and disease conditions.

**Highlights:**

- A flux model of energy metabolism integrating ^13^C labeling of metabolites and CO_2_
- Circulating nutrients have characteristic partition between oxidation and storage
- Circulating nutrients’ total cycling flux outweighs their total oxidation flux
- Cycling fluxes are extensively elevated in ob/ob but not in diet-induced obese mice

**Graphical Abstract:** 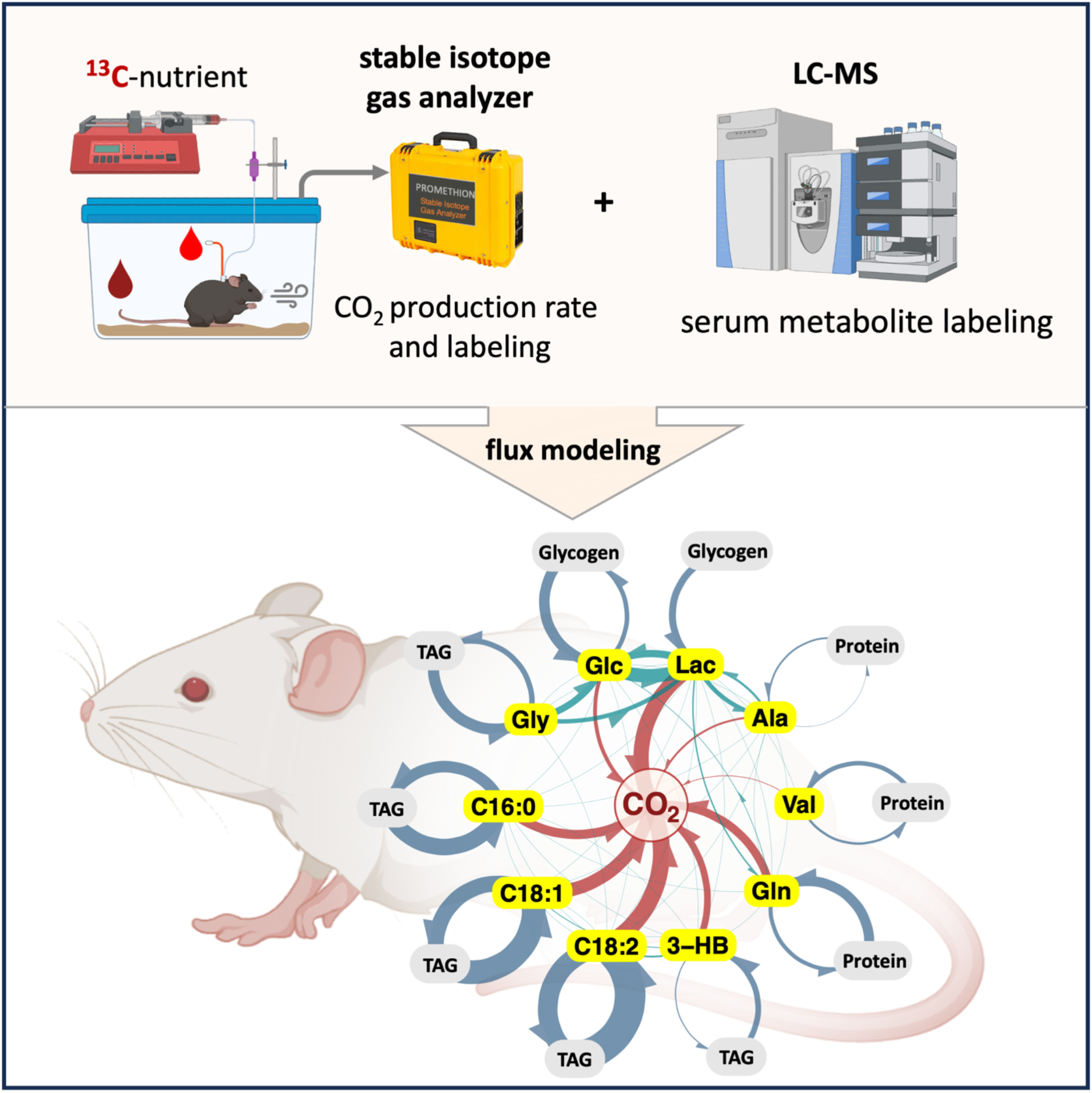

## INTRODUCTION

While our body is ultimately powered by nutrients in the food we eat, our tissues consume nutrients from the blood circulation for their energy needs, mostly by oxidizing them into CO_2_. The circulating nutrients are supplied from digestion of food or breakdown of tissue nutrient storages such as glycogen and triacylglycerol (TAG). Instead of following the simple one-way process of release from storage to oxidation by tissues, however, a circulating nutrient can be simultaneously recycled back to its tissue storages, thus forming metabolic cycles (**Figure 1A**). For example, circulating glucose can be released from and resynthesized into tissue glycogen at the same time (Shipley et al., 1967, 1970). Another well-known example is the triacylglycerol-fatty acid cycle, which involves the simultaneous release of non-esterified fatty acid (NEFA) from tissue triacylglycerol (TAG) into circulation and resynthesis of tissue TAG from circulating NEFA (Nye et al., 2008; Randle et al., 1963). In addition to the metabolic fates of oxidation and resynthesis into tissue storage, circulating nutrients also interconvert with each other, forming additional metabolic cycles. One prominent example is the Cori cycle, where circulating glucose is converted into circulating lactate, which is then converted back into circulating glucose through gluconeogenesis (Cori, 1931; Reichard et al., 1963). Thus, circulating nutrients are involved in a complex network of fluxes, including those in oxidation, inter-nutrient metabolic cycling, and nutrient-storage metabolic cycling (**Figure 1A)**. Despite extensive knowledge of some well-known examples, a holistic and quantitative view of these fluxes encompassing all major circulating nutrients is lacking, preventing a global view of how energy metabolism is functioning in health and diseases such as obesity.

**Figure 1.**
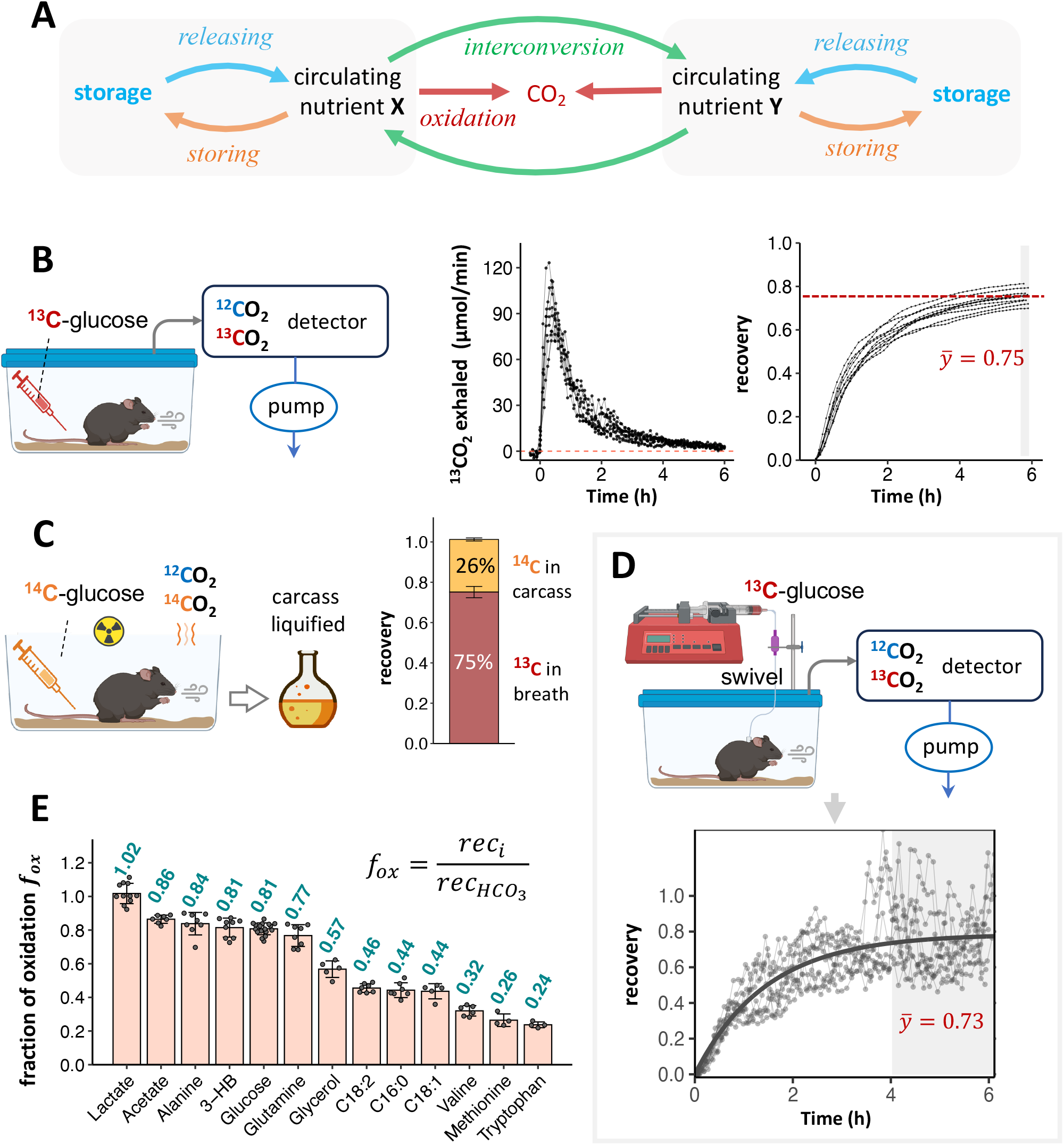
^13^CO_2_ Tracing Unveils Characteristic Oxidation and Storage Partition of Circulating Nutrients. **(A)** Schematic of the releasing, storing, oxidation, and interconversion of circulating nutrients. **(B)** Tracing of ^13^CO_2_ production after intravenous (i.v.) injection of a nonperturbative bolus dose of ^13^C-glucose. The ^13^CO_2_ production kinetics (middle) and accumulated ^13^C recovery (right) are shown. Each line represents data from one animal. n = 10. **(C)** Deposit of ^14^C radioactivity in the carcass 6h later after i.v. injection of ^14^C-glucose. n=3. **(D)** Tracing of ^13^CO_2_ production using nonperturbative i.v. infusion of ^13^C-glucose in conscious unrestrained animal. Shown are the experimental setup (top) and ^13^C recovery (bottom). Each thin line represents data from one animal and the thick line is fitted with formula y = a · (1 − e ^−b.t^). n = 10. **(E)** Values of fraction of oxidation (*f*_*ox*_) of 13 major circulating nutrients. Mean ± SD. n = 4 to 17 for individual nutrient, as noted in the scatterplot.

Isotope tracing is a powerful approach to determine metabolic fluxes. For a system (e.g., cells, animals) administered with isotope tracers, the isotope labeling of metabolites reflects the fluxes on the underlying metabolism (Fernández-García et al., 2020; Metallo, 2015). Modern analytical methods such as mass spectrometry (MS) and nuclear magnetic resonance (NMR) can generate rich isotope labeling information for many metabolites, allowing systematic flux calculations for multiple metabolic reactions within complex networks (Antoniewicz, 2018; Hasenour et al., 2020). The applications of isotope-based flux analysis has enabled many important discoveries in both in vitro and in vivo models (Befroy et al., 2014; Cappel et al., 2019).

In our previous work, with infusion of isotope-labeled nutrients to mice and mass spectrometry measurement of circulating metabolite labeling, we systematically determined the interconversion fluxes between the major circulating nutrients, revealing sustained metabolic cycling across diets (Hui et al., 2020).

In addition to the classic isotope analysis platform by MS and NMR, isotope-labeled CO_2_ tracing provides uniqiue insight of nutrient oxidization. While the tracing of radioactive ^14^CO_2_ was employed in the previous in vivo flux analysis, it has been done only to a few nutrients, such as glucose (Del Prato et al., 1993) and palmitate (Campbell et al., 1992), and simultaneous labeling measurement for multiple metabolites was restricted. The use of stable ^13^CO_2_ tracing (McCue and Welch, 2016), on the other hand, makes close integration with the MS measurement a possibility, creating a new yet uncharted revenue of in vivo flux quantification.

Here, we integrated isotope tracer infusion, MS, and ^13^CO_2_ gas analyzer measurement, and developed a fluxomics framework to calculate the oxidation, inter-nutrient and nutrient-storage fluxes for 10 major circulating nutrients in mice (**Figure S1**). The resulting quantitative flux model at the organism-level revealed many surprising features of mammalian energy metabolism. Of note, metabolic cycling flux is numerically more prominent than oxidation for circulating nutrients, despite the already enormous oxidative flux. To unravel the global changes of energy metabolism in obesity, we applied this framework on obesity models. Among our findings, compared with lean mice and on a per animal basis, the metabolic cycling fluxes of carbohydrate and fat nutrients were elevated by about 2-fold in the leptin-deficient obese mice while they remained largely the same in diet-induced obese mice. Overall, our work provides a framework for building an organism-level flux model of energy metabolism, a useful tool for metabolic research.

## RESULTS

### Quantifying the Fraction of Oxidation for a Circulating Nutrient

In general, the carbon atoms in a circulating nutrient either leave the body as the oxidation product CO_2_ or are recycled back into storages in the body (**Figure 1A**). To quantify these oxidation fluxes and storing fluxes of circulating nutrients, we first aimed to determine what fraction of a circulating nutrient is oxidized into CO_2_, or its fraction of oxidation (*f*_*ox*_). For this, using circulating glucose as an example, we intravenously administered a small amount of ^13^C-uniformly labeled glucose as a single bolus into modestly fasted mice, and monitored the ^13^CO_2_ production using a metabolic cage equipped with a ^13^CO_2_ analyzer (**Figure 1B**). The production flux of ^13^CO_2_ peaked at around 30 min, significantly dropped in another 2 hours, and tailed to approach background level within a total of 6 hours (**Figure 1B;** see calculation of ^13^CO_2_ flux in **Supplementary Note 1**). Integration of the ^13^CO_2_ production kinetic curve over the 6 hours gives the total ^13^CO_2_ production from the tracer administered. The recovery was then calculated as the ratio between the total production of ^13^CO_2_ (µmol) and the administered tracer dose (µmol ^13^C atoms). Glucose recovery was thus determined to be 0.75. This suggests that 25% of circulating glucose atoms were destined to storage pools, such as triglycerides, proteins, glycogen, and other macromolecules. To verify this number, we intravenously injected the radioactive tracer ^14^C-glucose into mouse, and measured the remaining radioactivity in the carcass 6 hours after administration. The carcass was digested into a solution, and the radioactivity was measured using a scintillation counter. We detected 26% of the administered dose in the carcass (**Figure 1C**), consistent with our ^13^CO_2_ measurement and a previous result (Tolbert et al., 1956).

To further confirm the glucose recovery result, instead of a bolus injection of ^13^C-glucose, we infused ^13^C-glucose into the mouse systemic circulation via a pre-implanted jugular vein catheter. Compared to the bolus injection, the constant infusion at a low rate reduces the amount of exogenous nutrient per minute administered to the animal and thus minimizes potential perturbation to metabolism. Here the recovery is the ratio of ^13^CO_2_ production rate (µmol/min) at steady state to the ^13^C-infusion rate (µmol ^13^C atoms/min). With this experiment, we measured glucose recovery to be 0.73 (**Figure 1D**), in close agreement with that measured with the bolus administration. We thus combined results from both methods and took the average as the recovery of glucose.

To obtain the *f*_*ox*_ value for glucose, we next took in account the fact that a portion of the ^13^CO_2_ generated from oxidation of ^13^C-glucose can undergo carboxylation and get incorporated into metabolite pools (such as urea and oxaloacetic acid) which are not recovered in the exhalation (Elia et al., 1992). For this, we intravenously infused ^13^C-labeled bicarbonate, and determined a recovery of 90% for bicarbonate, indicating that 90% of produced CO_2_ from substrate oxidation actually leaves the body as exhaled CO_2_. We thus took the ratio of glucose recovery to bicarbonate recovery as the glucose *f*_*ox*_, and obtained a value of 0.81 (**Figure 1E**). This value reflects that ∼80% of circulating glucose is oxidized, and ∼20% is recycled back to storage pools independent of the CO_2_ carboxylation pathway.

### Distinct *f*_*ox*_Values for Different Circulating Nutrients

In addition to glucose, we went on to quantify the *f*_*ox*_ for 12 other circulating nutrients (**Figures 1E, S1B-E**). For most circulating nutrients investigated, the *f*_*ox*_ by both bolus injection and infusion methods are generally consistent (**Figure S1D**), and we pooled the data from both methods. Glycerol, however, was 30% more oxidized when administered as a bolus than by infusion, suggesting significant perturbation to glycerol metabolism by the bolus administration. Given the negligible perturbation by infusion, we used infusion-based *f*_*ox*_ for glycerol for further analyses.

Among all 13 circulating nutrients, lactate had the highest *f*_*ox*_ of around 100%, significantly higher than glucose *f*_*ox*_ of 81% (**Figure 1E**). This result further substantiates circulating lactate’s role as an energy source for tissues (Hui et al., 2017). The ketone body 3-hydroxybutyrate and acetate also have high *f*_*ox*_ of 81% and 86%, respectively, indicating their main use as energy sources in the body. Non-esterified fatty acids (NEFA) occupy the lower end of the *f*_*ox*_ spectrum. Palmitate (C16:O), oleate (C18:1), and linoleate (C18:2), the most abundant NEFAs in circulation, have *f*_*ox*_ values of around 45%, suggesting prominent fluxes from circulating NEFAs back to tissue triglycerides (Reshef et al., 2003). The branched-chain amino acid (BCAA) valine has a *f*_*ox*_ of only 32%, similar to that of BCAA leucine (Tolbert et al., 1956). Together with *f*_*ox*_ of 24∼26% for the essential amino acids methionine and tryptophan, this result reveals that the majority of proteolysis-released circulating amino acids are recycled back into proteins. In contrast, circulating glutamine and alanine have high *f*_*ox*_ of 77% and 84%, respectively, indicating their unique roles in energy metabolism among amino acids. Thus, *f*_*ox*_ of a circulating nutrient reveals its main role in mammalian metabolism and is a fundamental property of circulating nutrients.

In the following flux analysis, we focused on 10 nutrients in **Figure 1E**, which were the top 9 circulatory nutrients (Hui et al., 2017) and valine. Albeit a small circulatory turnover flux, valine was included as a reporter of protein metabolism. The other 9 nutrients are glucose, lactate, glycerol, alanine, glutamine, 3-HB, C16:0, C18:1, and C18:2.

### Quantifying Oxidation and Storing Fluxes of Circulating Nutrients

Based on the *f*_*ox*_ measurement, we next sought to quantify the fluxes associated with oxidation and storing destinies of circulating nutrients. At the steady state of ^13^C-tracer infusion, we sampled the blood and measured the ^13^C-labeling of the infused nutrient using mass spectrometry (**Figure 2A**). The serum labeling of the infused nutrient (*L*) reflects the dilution of the infusate (at infusion rate *I*) by the endogenous production flux *R*_*a*_ of the nutrient, with 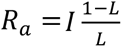. The *R*_*a*_ is an important quantity to reflect a nutrient’s circulatory turnover, and can be significantly changed at different physiological conditions, such as under hyperinsulinemia (**Figure S2A**) and beta-adrenergic agonism (**Figure S2B**). Since *f*_*ox*_ and 1 − *f*_*ox*_ are the fraction of *R*_*a*_ that is fated to oxidation and storing, respectively (**Figure 2B**), the oxidation and storing fluxes of a circulating nutrient, denoted as *J*_*ox*_ and *J*_*S*_, respectively, can be calculated (see derivation in **Supplementary note 2**) as

**Figure 2.**
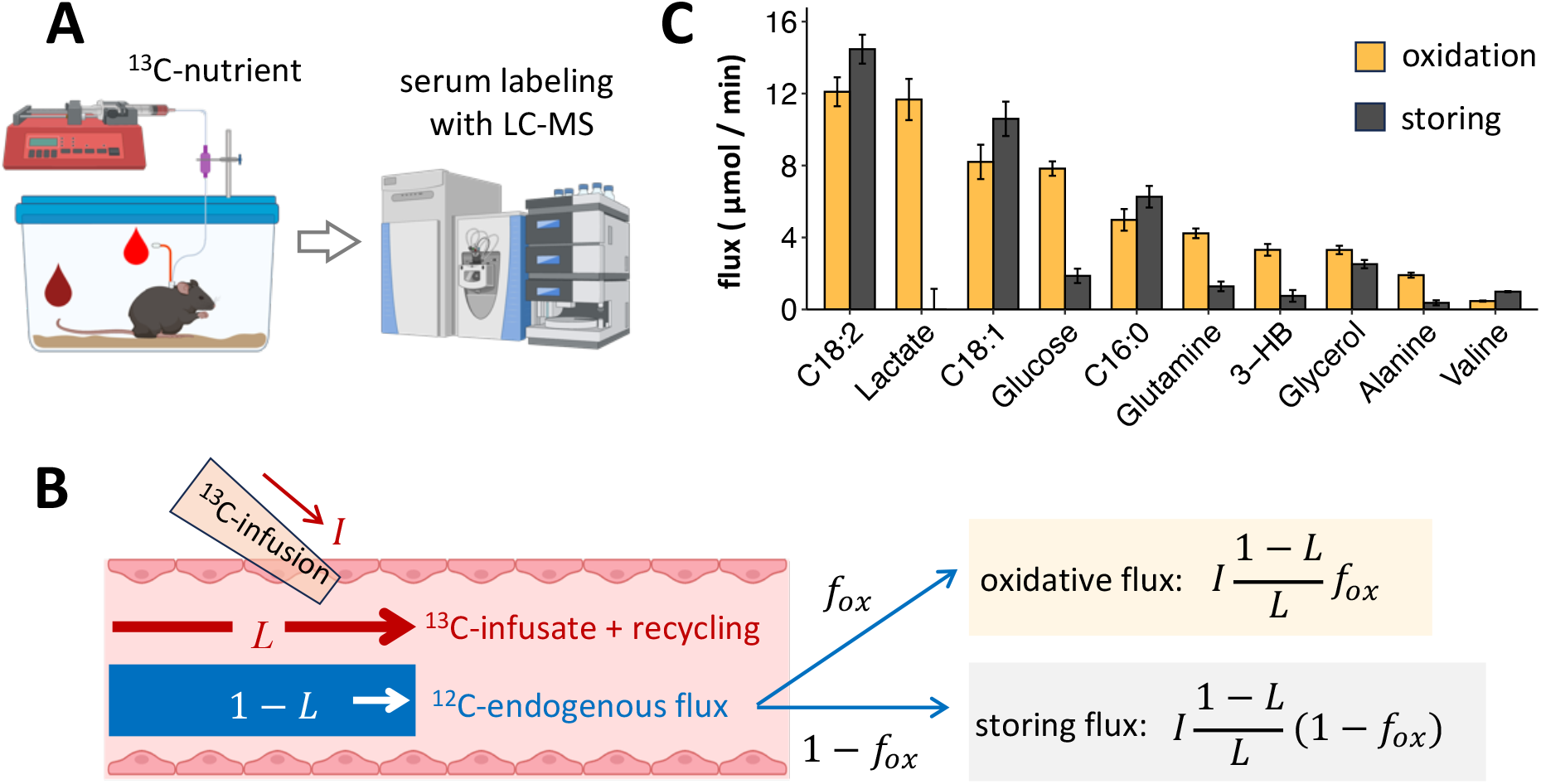
Quantification of the Oxidation and Storing Fluxes of Circulating Nutrients. **(A)** Schematic showing infusion of ^13^C-nutrient to mice and analysis of the serum labeling with LC-MS. Blood was collected from arterial blood via pre-implanted catheter and/or by tail snip. **(B)** Illustration of the formula for calculating oxidation and storing fluxes of circulating nutrients. *I*, infusion rate of ^13^C-nutrient; *L*, serum labeling of the infusate; and *f*_*ox*_, fraction of oxidation, quantified in Figure 1. **(C)** Oxidation and storing fluxes of major circulating nutrients. Mean ± SEM. For individual nutrient, for their labeling in serum, n = 4∼15 (see Figure S3A); for *f*_*ox*_, n = 4 to 17 (Figure 1E); the same replication applies to all figures computed from these two sets of parameters in Figures 2-5 and related supporting figures. Fluxes normalized by body weight refers to Figure S2C.

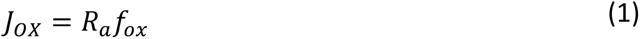

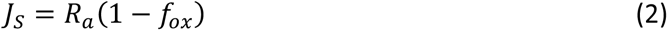

The absolute flux values reveal quantitatively the magnitude of each nutrient as an oxidation substrate or storage precursor (**Figure 2C**). For example, though linoleic acid has lower *f*_*ox*_ than glucose (**Figure 1E**), it has higher oxidation flux. Valine has the largest storing fraction (1 − *f*_*ox*_), but the associated storing flux is among the smallest.

### Quantifying *Direct* Oxidation and Storing Fluxes of Circulating Nutrients

The oxidation flux for a circulating nutrient quantified above describes the nutrient’s ultimate disposal into CO_2_. It includes both direct and indirect oxidation fluxes of the nutrient. The indirect oxidation refers to the process that a circulating nutrient is first converted into other circulating nutrients before being oxidized. For example, an important indirect oxidation route for circulating glucose is via circulating lactate. In like manner, the storing flux also comprises both the direct and indirect ways. To gain further insights on the sources and sinks of circulating nutrients, we next sought to quantify the *direct* fate of a circulating nutrient to oxidation and storage.

We denote the direct oxidation flux from a circulating nutrient *j* as *O*_*j*_, which we aim to calculate. Now consider the steady state of a nonperturbative infusion of a ^13^C-labeled nutrient, e.g., ^13^C-glucose. The ^13^CO_2_ production flux from the animal is given by *If*_*ox*_, where *I* is the infusion rate and *f*_*ox*_ is the glucose fraction of oxidation. As the ^13^CO_2_ production is also equal to the sum of labeled direct oxidation fluxes from all circulating nutrients, we have

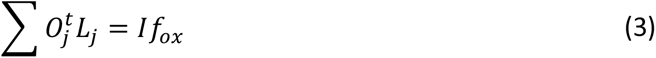

where 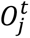 is the direct oxidation flux (including both ^13^C and ^12^C) of nutrient *j*, and *L*_*j*_ the labeled fraction of the nutrient (illustrated in **Figure 3A**). As the endogenous unlabeled oxidation flux *O*_*j*_ accounts for 1 – *L*_*j*_ fraction of 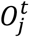, or 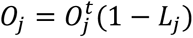, plugging it into the above equation gives the expression with the endogenous oxidation flux *O*_*j*_ as the only unknown

**Figure 3.**
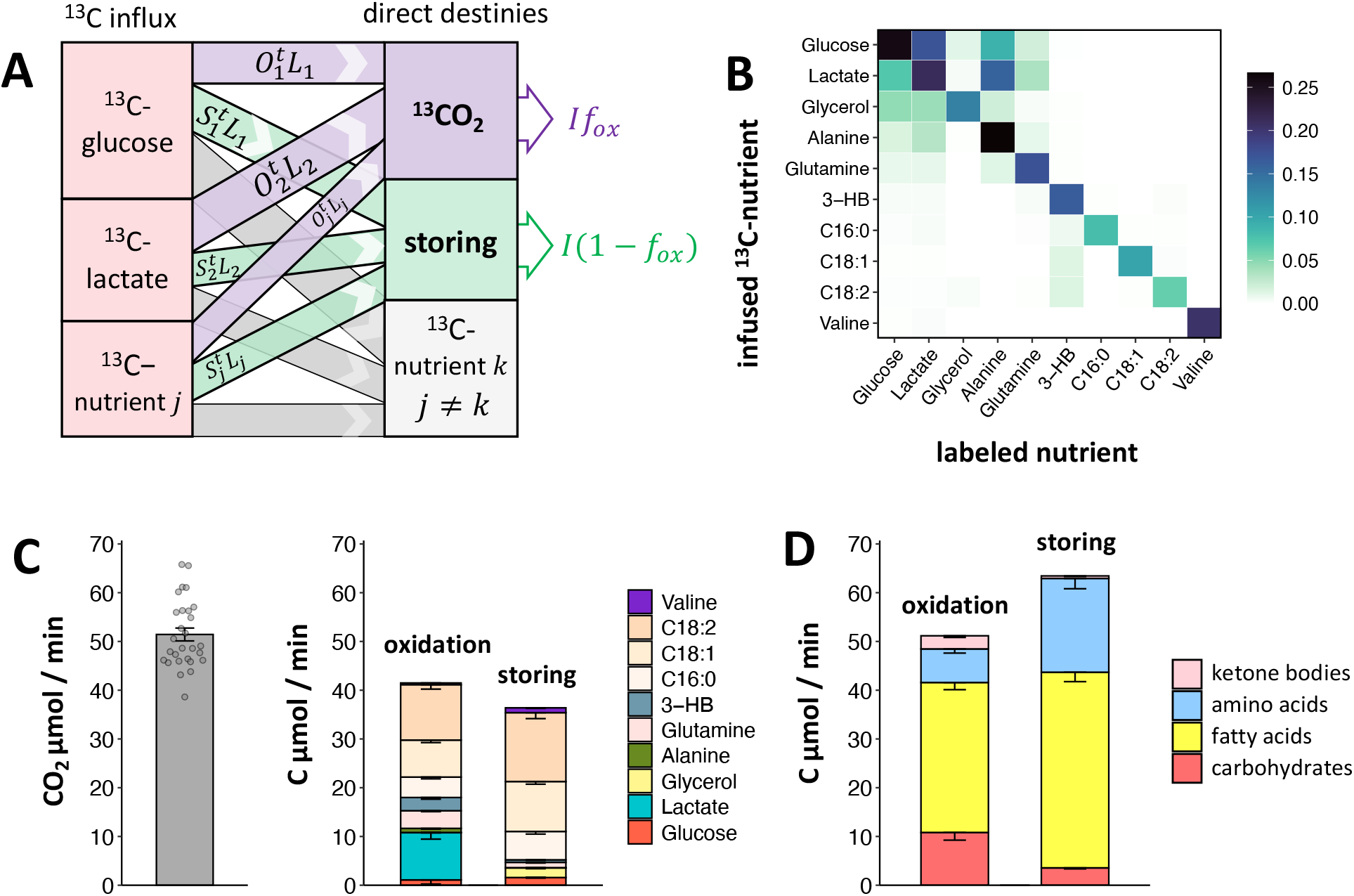
Quantification of *Direct* Oxidation and Storing Fluxes of Circulating Nutrients. **(A)** Illustration for calculating direct oxidation and storing fluxes of circulating glucose. At the isotopic steady state of ^13^C-glucose infusion, the total ^13^CO_2_ production (oxidation) and storing flux of the ^13^C-infusate are depicted as the sum of the direct oxidation and storing fluxes from all labeled circulating nutrients, respectively. 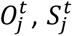 and *L*_*j*_ are respectively the oxidation flux, storing flux (superscript *t* for total, including both ^12^C and ^13^C), and ^13^C labeling of circulating nutrient *j*; *f*_*ox*_, fraction of oxidation. **(B)** Labeling of circulating nutrients under the infusion of different ^13^C-nutrients. n = 4 to 15 for individual nutrient (see Figure S3A). **(C)** Total CO_2_ production (left, n = 28), in comparison with the direct oxidation and storing fluxes of the major circulating nutrients covered in this study (right). **(D)** Direct oxidation and storing fluxes of all major circulating nutrients, including carbohydrates (glucose, lactate, and glycerol), fatty acids, protein-originated amino acids, and ketone bodies. All data is presented in mean ± SEM. For panels (C-D), normalized fluxes by body weight refers to Figure S3C-D.

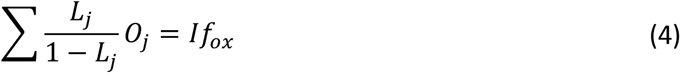

This equation holds under the infusion of each different ^13^C-labeled nutrient. As such, the infusion of 10 different ^13^C-nutrients unveils the inter-labeling pattern among these nutrients (**Figures 3B** and **S3A**), and allows the construction of a linear system of 10 equations of

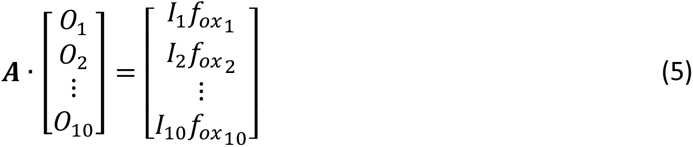

where ***A*** is a 10×10 matrix, with element 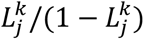 in the *k*^*th*^ row and *j*^*th*^ column, and 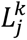 representing the labeling of *j*^*th*^ circulating nutrient under the infusion of the *k*^*th*^ tracer. Solving these linear equations yields the direct oxidation fluxes *O*_*j*_ of the circulating nutrients (**Figure 3C**). The direct storing fluxes *S*_*j*_ can be calculated in the same manner (**Figure 3C**; see more details in derivation in **Suppl. Note 3**).

### Direct Contribution of Circulating Nutrients to Total Energy Expenditure

The direct oxidation fluxes form an organism-level view of burning circulating nutrients for energy (**Figure 3**). Compared with the total CO_2_ exhaled by the animal measured directly using a metabolic cage, the 10 nutrients alone accounted for about 80% of the total CO_2_ production, confirming that these nutrients represent the top circulating nutrients. Glucose, while being the most famous energy nutrient, has rather minimal direct oxidation flux, accounting for only marginal fraction of total energy expenditure (TEE) (**Figure 3C**). Instead, circulating lactate, mostly derived from glucose, is the dominant carbohydrate that directly fuels oxidation (Hui et al., 2017), responsible for about 20% of TEE. Glutamine, as the single most important amino acid fuel, contributes to 7% of TEE, while alanine and valine contribute 1.5% and 0.7%, respectively. As the oxidative disposal of essential amino acid reflects the use of proteins for energy expenditure, based on the content of valine (5%) in muscle proteins (Gorissen and Phillips, 2018), the direct oxidation of protein-originated amino acids can be estimated to provide approximately 14% of TEE (Fearon et al., 1988; Matthews et al., 1980). The ketone body 3-HB contributes to 5% of TEE. Due to the rapid equilibrium between 3-HB and acetoacetate, this quantity reflects the contribution of the total ketone bodies (Keller et al., 1981). The three NEFA species, palmitate, oleate, and linoleate, are among the most prominent fuels, contributing to 8%, 15% and 22% of TEE, respectively, or 45% in sum. As these top 3 NEFAs account for about 75% of total NEFA in circulation (**Figure S3B**), approximately 60% of TEE is supported by the total NEFA in circulation. Glycerol is uniquely the only nutrient that is not directly engaged in oxidation (Hui et al., 2020). All nutrients considered together, using both direct measurement and empirical extrapolation, close to 100% of TEE can be quantitatively accounted for (**Figure 3D**, showing the direct TEE contribution from carbohydrate nutrients (glucose, lactate, and glycerol), NEFAs, protein, and ketone bodies).

### Recycling to Storage Is a Highly Prominent Fate of Circulating Nutrients

The direct storing fluxes revealed the composition of nutrients for resynthesizing fuel storages. The total NEFAs present a dominating 60% of the total storing flux (**Figure 3D**). Amino acids also have prominent storing fluxes. Using valine as a representative of protein-originated amino acids, the total amino acids storing flux accounts for 30% of the total storing flux. Carbohydrates have a minor part (6%) in the total storing flux, with lactate having zero direct storing flux. This again highlights lactate’s role as a fuel metabolite. Most strikingly, all circulating nutrients considered together, the total storing flux is 23% higher than that of oxidation (**Figure 3D**), vividly demonstrating the enormity of futile cycling fluxes of circulating nutrients.

### Quantifying Inter-converting Fluxes between Circulating Nutrients and their Releasing Fluxes from Storages

We have so far focused on oxidation and storing as destinies for circulating nutrients. The storages however also serve as sources for circulating nutrients. To work toward a full view of metabolism of circulating nutrients, here we attempted to quantify the production fluxes of the circulating nutrients from the storage pools. In the same calculation, we also obtained the interconverting fluxes between circulating nutrients, which was a main result in our previous work (Hui, et al. 2020).

To illustrate the calculation, we use glucose as an example and quantify its production fluxes, including the storage releasing flux *F*_*S*_, i.e., glycogenolysis, and the gluconeogenic flux *Fj* from circulating nutrient *j* (**Figure 4A**). Now consider the steady state during a nonperturbative ^13^C-glucose infusion with an infusion rate *I*. The total glucose production flux (*P*^*total*^) at steady state is equal to summation of all incoming fluxes from different sources to glucose, given by

**Figure 4.**
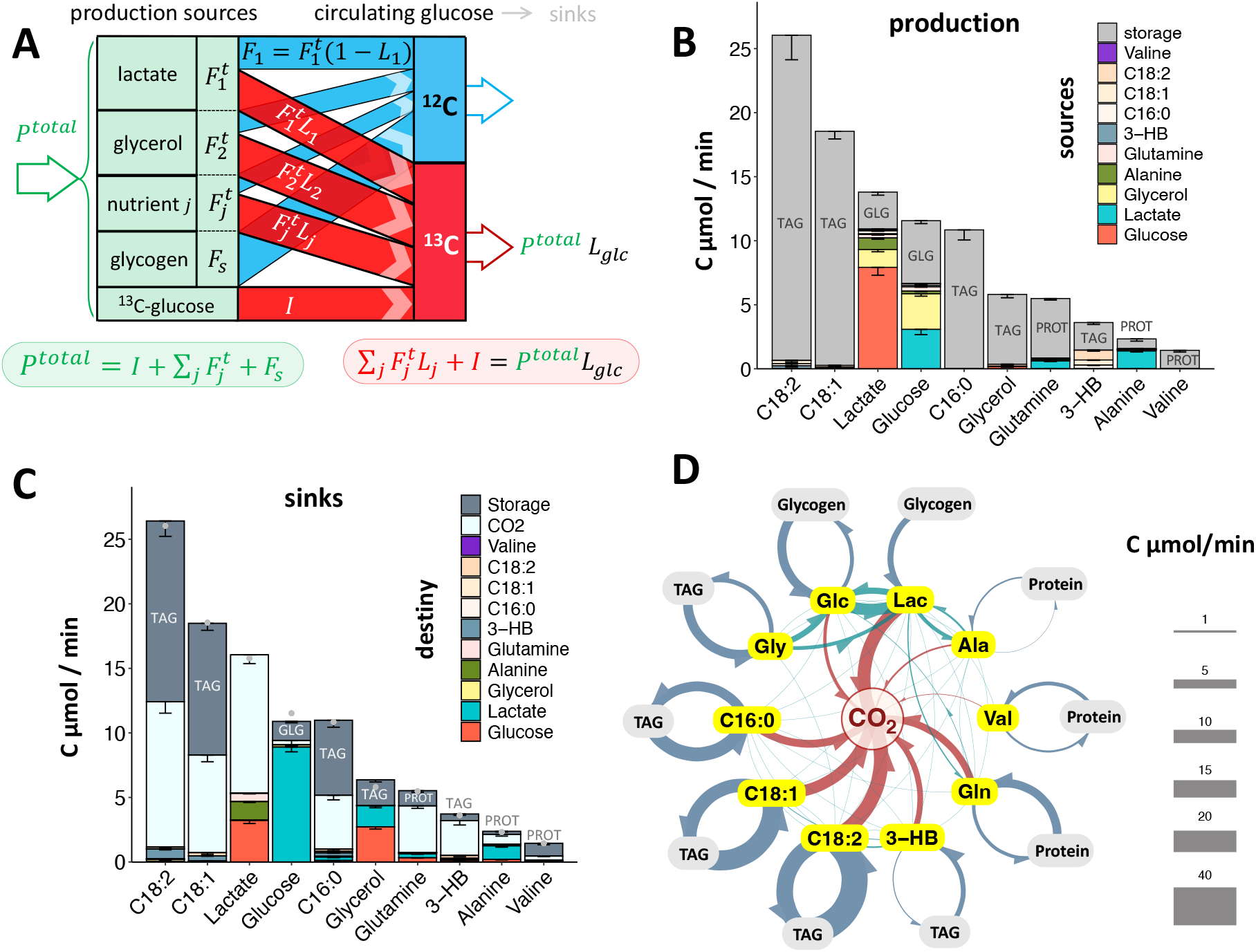
Quantifying Storage Releasing Flux and Interconverting Fluxes for Circulating Nutrients, and Integrating them with Oxidation and Storing Fluxes into a Holistic Flux Model of Circulating Nutrients. **(A)** Illustration for calculating the storage releasing flux (glycogenolysis) and gluconeogenic fluxes to produce circulating glucose. *I*, the ^13^C-glcose infusion rate; *L*_*glc*_, the labeling of glucose; *L*_*j*_, the labeling of nutrient *j*; 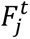, the production flux from nutrient *j* to glucose, superscript *t* for total (including both ^12^C and ^13^C); *F*_*S*_, the releasing flux from storage of glycogen. **(B)** The production fluxes of circulating nutrients from storage and other circulating nutrients. TAG, triglyceride; GLG, glycogen, PROT, protein. **(C)** Sink fluxes of circulating nutrients, fated to oxidation, storage, and other circulating nutrients. The dot shows the total production flux as quantified in Figure 4B. For panels (B) and (C), fluxes normalized by body mass refers to Figure S4 A-B. **(D)** An organism-level metabolic map of circulating nutrients. All flux arrows are drawn in counter-clockwise direction. All data is presented in mean ± SEM.

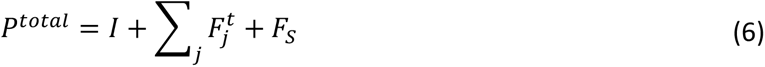

where 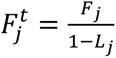 with *Lj* the labeling of nutrient *j*. Note that 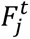 is the total flux from nutrient *j* to glucose including both endogenous unlabeled flux *F*_*j*_ and labeled flux 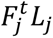. The mass balance of the labeled glucose pool (with the labeling denoted as *L*_*glc*_) requires

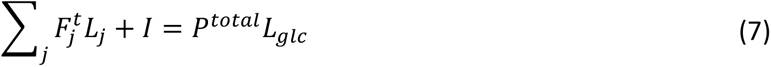

where the left-hand-side is the incoming ^13^C flux to glucose from different sources, and the right-hand-side the outgoing labeled glucose flux. Eq. (6) and the expression for 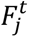 can be plugged into Eq. (7) to form an equation with *F*_*j*_’s and *F*_*S*_ as the only unknowns. Such an equation holds under the infusion of other ^13^C-nutrients, but with *I* = 0. With infusions of ^13^C glucose and 9 other ^13^C-labeled circulating nutrients, a linear system of 10 equations can be then constructed for the 10 unknowns including the 9 *F*_*j*_ ‘s and *F*_*S*_, in like manner as Eq. (4) (see Methods). Solving the equations yielded the storage releasing flux, and the conversion fluxes from the 9 circulating nutrients to produce circulating glucose. In similar fashion, the releasing fluxes and inter-converting fluxes can be calculated for other circulating nutrients using the same computational framework. More derivation details can be found in **Suppl. Note 4**.

### Sources of Circulating Nutrients

The storage releasing flux and inter-converting flux results calculated above provide a quantitative view of the direct sources for each of the 10 circulating nutrients (**Figure 4B**). Two types of circulating nutrients can be seen, nutrients directly sourced from the storages, and those derived from both storages and other circulating nutrients. NEFA and glycerol are almost exclusively released from the TAG store, and valine and glutamine from the protein store. In contrast, glucose, lactate, alanine, and 3-HB are additionally sourced from other circulating nutrients. For circulating glucose, about 40% is released from glycogen, while the remaining majority is produced by gluconeogenesis, primarily from lactate and glycerol (each about 25%). For circulating lactate, about 20% is derived from intracellular glycogen breakdown and subsequent glycolysis, in line with that measured by pulse-chase tracing of glycogen (TeSlaa et al., 2021), while the remaining mostly comes from circulating glucose (60%) and glycerol (10%). As glycerol is derived from TAG, this collectively shows that glycogen and TAG but not protein are the major ultimate sources of circulating glucose (Wang et al., 2020). We further quantified such ultimate contribution fraction from various storages to circulating nutrients (**Figure S4C**, see derivation in **Suppl. Note 5**). Interestingly, while amino acids are typically released from the protein as the principal source, alanine has its carbon origin roughly evenly from protein, glycogen, and TAG.

### Sinks for Circulating Nutrients

Together with the direct oxidation and storing fluxes calculated earlier (**Figure 3C**), the inter-conversion fluxes of circulating nutrients constitute the direct sinks of circulating nutrients (**Figure 4C**). This integrated view offers interesting insights. For example, though glutamine and alanine, are known to be gluconeogenic, their direct conversion to lactate is more predominant than to glucose. In addition, these amino acids are directly oxidized with higher fluxes than otherwise relying on the lactate-genic pathway for energy supply.

### A Flux Model Incorporating All Quantified Fluxes for Circulating Nutrients

A further inclusive combination of the production and consumption fluxes resulted in a comprehensive metabolic flux model of circulating nutrients (**Figure 4D**). In addition to the prominent oxidative fluxes for energy generation, this model vividly highlights two types of cycling of the circulating nutrients. One type involves the cycling between the circulating nutrients and the storages, and a second cycling exists between the circulating nutrients. These cycles render a picture of vibrant carbon exchanges among nutrients in circulation and tissues. Many of the futile cycles proceed at the cost of ATP. We estimated ATP wasted in the Cori cycle, TAG-NEFA cycling, and protein turnover, the top three cycles with the highest carbon flux (Methods). Together, these cycles wasted remarkably around 10% of the total ATP generated (**Figure S4D**).

### Fluxes Are Generally Unperturbed in Diet-Induced Obese Mice

To further demonstrate the utility of the systems-level flux modeling, we applied the framework to two obese mouse models, high fat diet-induced obese mice (HFD), and leptin-deficient ob/ob mice (**Figures 5-6, S5**). Unexpectedly, despite being 60% heavier than the wild-type control mice on chow diet, the HFD mice showed generally moderate changes in *f*_*ox*_ and fluxes (**Figures 5A-D**). The most striking change is the significantly increased production of circulating glycerol from glucose. While it was elevated by 2-fold in ob/ob mice, it was more impressively activated by 4.5-fold in the HFD group (**Figures 5B, S6, S7A**). The production of glycerol from glucose may be derived from the recently discovered glycerol shunt pathway (Mugabo et al., 2016; Possik et al., 2023), or from the spillover of the glycerol backbone in the very-low-density lipoprotein (VLDL)-TAG during lipase hydrolysis, where the backbone is synthesized from blood glucose in the liver (Jensen et al., 2001).

**Figure 5.**
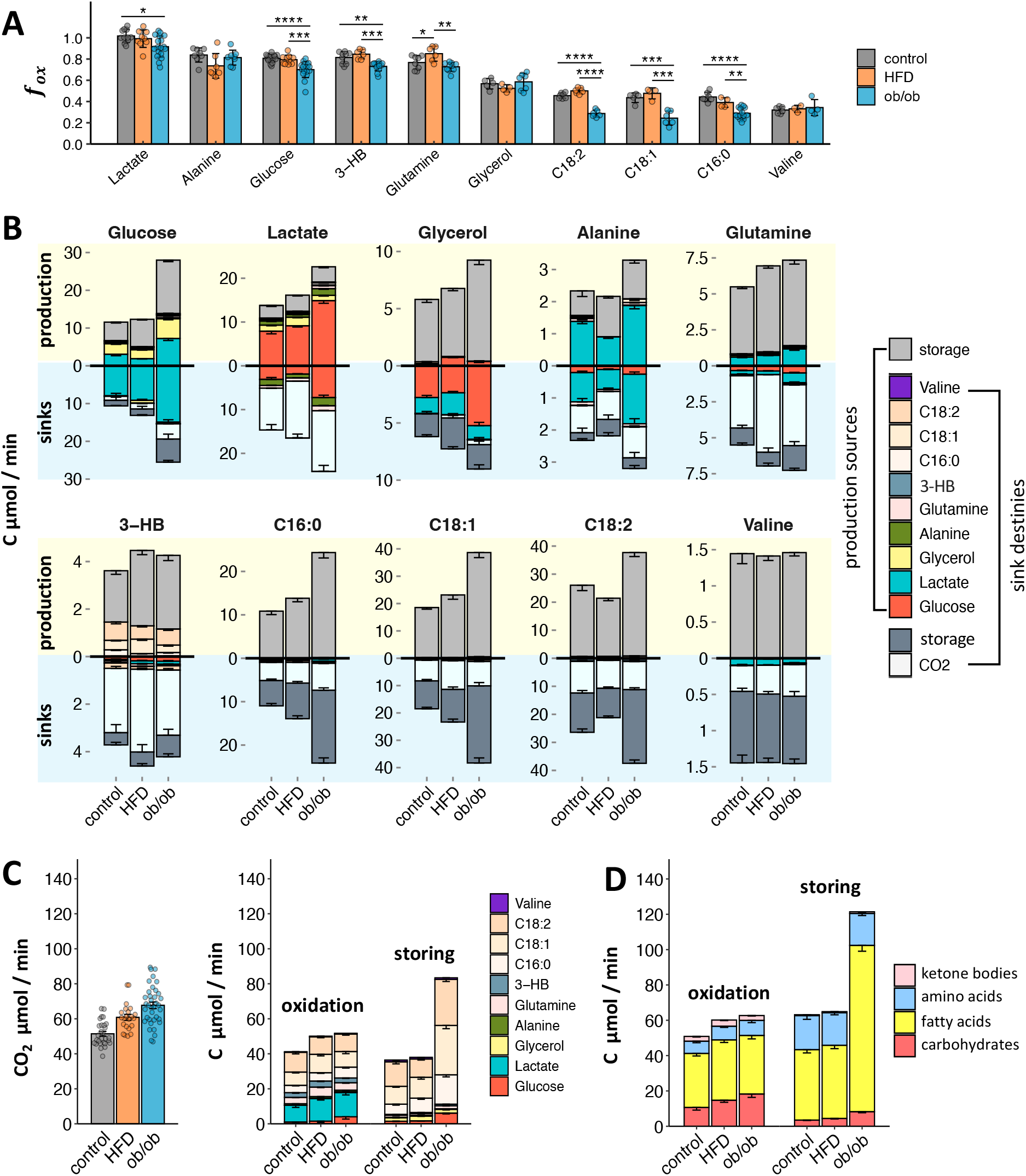
Systematic Flux analysis of Circulating Nutrients in High Fat Diet-Fed (HFD) and ob/ob Mice. **(A)** Fraction of oxidation (*f*_*ox*_) of circulating nutrients in chow-fed wild type control mice, and the obese mice. Mean ± SD. **(B)** Production and sink fluxes of circulating nutrients in the control and obese mice. **(C)** Total CO_2_ production (left), and the direct oxidation and storing fluxes of circulating nutrients (right) in the control and obese mice. **(D)** Direct oxidation and storing fluxes of all major circulating nutrients, including carbohydrates (glucose, lactate, and glycerol), fatty acids, protein-originated amino acids, and ketone bodies. Fluxes normalized by lean or body mass refers to Figure S6B. All data is presented as mean ± SEM unless otherwise specified. \**p* < 0.05, \*\**p* < 0.01, \*\*\**p*<0.001, \*\*\*\**p*<0.0001 by Student’s *t* test after Bonferroni adjustment. Regarding individual nutrient and phenotype, for *f*_*ox*_, n = 3 to 19 (see scatterplot in panel A); their labeling in serum, n = 4 to 27 (see Figure S6A). The same replication also applies to the related supporting figures.

**Figure 6.**
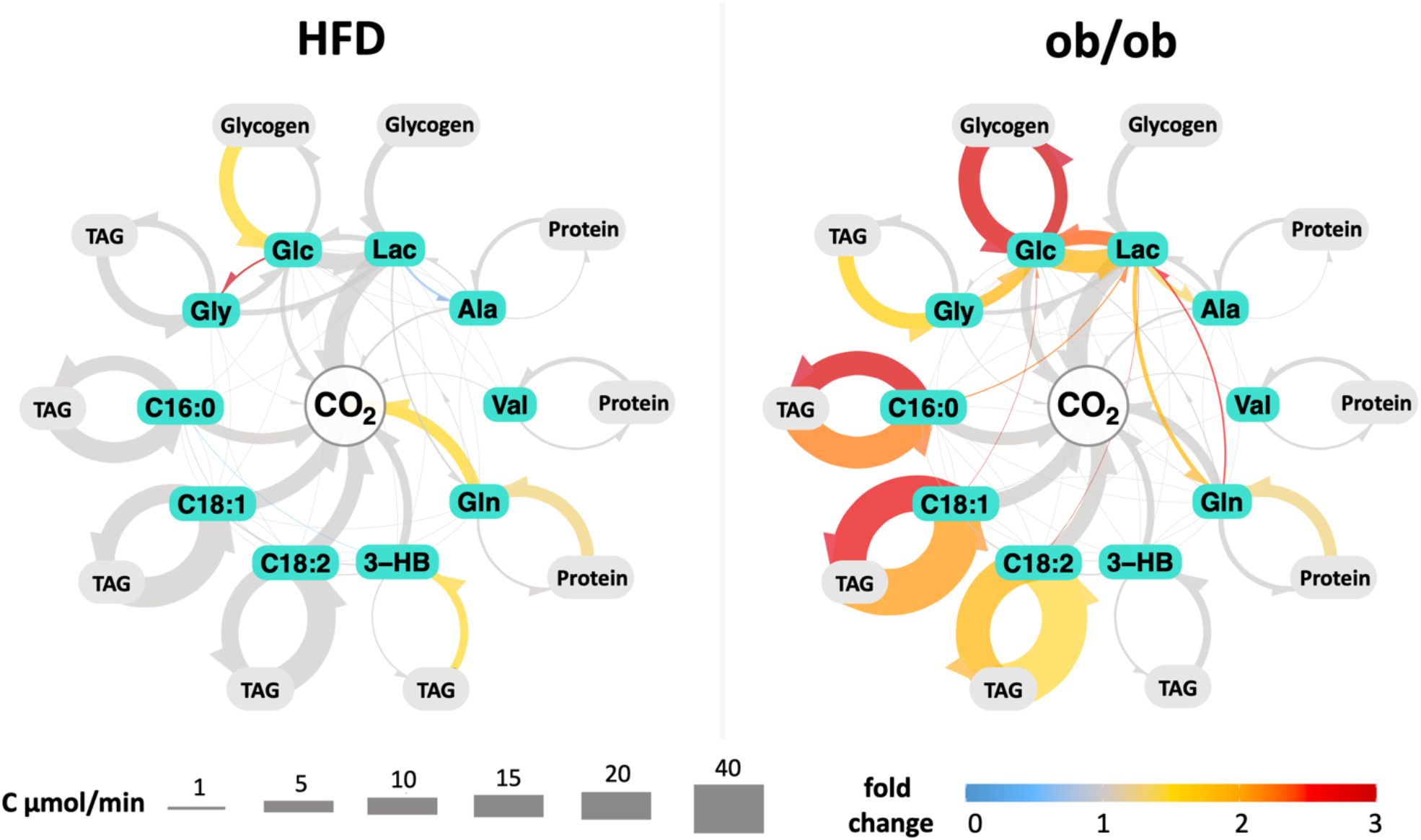
Flux Models of Obese Mice Reveals Elevated Metabolic Cycling in ob/ob Mice, but Not in High Fat Diet-Fed Mice. Fluxes in obese mice that are statistically significant (Bonferroni-adjusted *p* value < 0.05) from the control mice are highlighted in color, with fold change mapped to the color scale. All flux arrows are drawn in counter-clockwise direction.

### Extensive Elevation of Metabolic Cycling in ob/ob Mice

In contrast to the HFD mice, ob/ob mice showed major changes in energy metabolism. While *f*_*ox*_ was moderately decreased for many nutrients, it was most prominently decreased by 2-fold in circulating NEFAs (**Figures 5A**). This was accompanied by a 2-fold increase on average in the NEFAs’ circulatory turnover flux (equal to total consumption flux and total production flux) (Turner et al., 2005), resulting in their unchanged oxidation flux, but a 2.3-fold elevation in the recycling flux (**Figures 5B-D**). Intracellular reesterification nonetheless remained unchanged (**Figure S8A**). The circulatory fluxes of both glucose and lactate were also increased by around 2-fold (Turner et al., 2005), with production from different sources roughly proportionally scaled up. This led to a 2.3-fold increase in the oxidation of the carbohydrate nutrients (coupled with higher respiratory exchange ratio, **Figure S5D**), and over 2-fold increase in the fluxes of glucose recycling to storage and Cori cycle. While the increased metabolic fluxes in ob/ob mice may be attributed to a larger body mass, this effect can be statistically teased out using the generalized linear model approach (Mina et al., 2018). Statistical adjustment for body weight suggests that the augmented metabolic fluxes in ob/ob mice are more likely a result of metabolic rewiring inherent to the phenotype (**Figure S7B**). In addition, the flux distinction retains even when the ob/ob mice are compared to body-weight matched HFD mice (**Figure S7B**). Interestingly, as gauged by the essential amino acid valine, the global protein oxidation and futile recycling remains unchanged. All nutrients considered, the futile storing flux was 60% higher than their oxidation flux. The total ATP wasted in the top 3 futile cycles, TAG-NEFA extracellular recycling, Cori cycle and protein turnover, was 34% higher in ob/ob mice than the chow-fed control mice (**Figure S8B**).

## DISCUSSION

The present flux framework is a major advancement over our previous work (Hui et al., 2020), where we systematically quantified the inter-converting fluxes between circulating nutrients using isotope labeling data of serum metabolites. Here, by integrating ^13^CO_2_ tracing to the classic mass spectrometry-based isotope tracing framework, our framework further outputs the oxidation, storage and releasing fluxes of circulating nutrients. Together with the inter-converting fluxes, these fluxes form a full account of sources and sinks for circulating nutrients at the organism level, resulting in a coherent flux model of energy metabolism. Analogous to a map showing traffic along major highways in a city, our flux model captures the top fluxes in an animal body, providing an overview of energy metabolism landscape for the animal. Such a flux model for a disease state reveals unbiasedly the major alterations in energy metabolism for that disease, setting the stage for further investigations.

Our flux model revealed extensive metabolic cycling in the fasted state, with higher total flux than total oxidation flux. During fasting, the animal body is in a net catabolic state, with nutrient storages such as glycogen, TAG, and proteins being actively broken down to replenish circulating nutrients for oxidation into CO_2_. Yet, recycling back into the storage pools is a quantitatively more significant fate for circulating nutrients. What is the function of the extensive cycling? One hypothesis is that it is to produce heat, to maintain or elevate body temperature (Brownstein et al., 2022; Oeckl et al., 2022; She et al., 2007; Wolfe et al., 1987). We estimated substantial energy used by these futile cycles, about 10% total ATP generated. Another probable hypothesis is that metabolic cycling is to increase the sensitivity of metabolic regulation by enhancing the response magnitude for changes in the stimulus (e.g., substrate concentration)(Newsholme, 1978). We further revealed distinct partition between oxidation and cycling fluxes for different circulating nutrients, e.g., about 80% of circulating glucose is burned, while it is only ∼40% for a fatty acid, and ∼30% for an essential amino acid. The larger extent of metabolic cycling in fatty acids and amino acids may thus reflect their relatively less hormonal control than glucose. These and other hypotheses regarding the underlying principles of the presence of extensive cycling flux and of distinct *f*_*ox*_ values across nutrients deserve further investigations.

Our comprehensive flux results allow for comparison of fluxes between metabolic processes and can expose flux mismatches and thus knowledge gaps. This way, we identified an important gap in our understanding of the TAG-NEFA cycle, specifically regarding the sources of glycerol backbone for resynthesis of TAG. The flux through the TAG-NEFA cycle constitutes the major fraction of metabolic cycling, indicating active re-esterification of circulating NEFAs in adipose tissues (Kalderon et al., 2000). Glucose is classically viewed as the main source for glycerol backbone in TAG synthesis in adipose tissues (Sharma et al., 2024). However, even if all the glucose storing flux is diverted to adipose tissues for producing glycerol backbone, it would be insufficient to meet the demand (**Figure S8C**). Though lactate has been proposed as a supply of the glycerol carbons in the adipose tissue via the glyceroneogenesis pathway (Hanson et al., 2006), our data shows that circulating lactate has negligible flux fated to storages. Instead, our quantification shows that circulating glycerol has significant flux to storage, numerically and hypothetically fulfilling most of the needed glycerol backbone for TAG-resynthesis (**Figure S8C**). However, on the other hand, adipose tissue lacks glycerol kinase, and cannot sequester circulating glycerol directly. Therefore, the missing carbon source to synthesize TAG-glycerol backbone remains an important topic to be further investigated.

The flux results of HFD and ob/ob obesity models demonstrated strikingly distinct energy metabolism between them. Energy metabolic fluxes were mostly maintained in the HFD mice while there was extensive elevation of cycling fluxes in the ob/ob mice. While body weight differences can be a contributing factor, analysis of covariance (ANCOVA) with adjusted body weight unveils that the flux differences are more likely a result of intrinsic distinctions between phenotypes. Distinct fluxes in body weight-matched HFD and ob/ob group further supports this viewpoint. It is possible that the HFD mice resemble the “healthy obesity” group in human population, who are metabolically healthy despite being obese. On the other hand, the ob/ob mice lie in a totally different part of the energy metabolism space, possibly mirroring their departure from the state of metabolic health. Thus, instead of the commonplace view that ob/ob mice are an “artificial” obesity model, HFD mice and ob/ob mice may model two states along the progression of obesity.

The elevated glucose production and consumption fluxes in ob/ob mice inform on the cause of hyperglycemia. In general, an increase of a metabolite pool could result from either a blockage in the consumption reactions or an elevation in the production reactions. As production and consumption balance each other in the new steady state, the flux through the pool can be either reduced (in the consumption blockage case) or increased (in the production elevation case) compared to the unperturbed state. By this argument, the increased flux through the glucose pool in the ob/ob mice indicates elevated glucose production as the cause of hyperglycemia, implicating liver insulin resistance instead of muscle insulin resistance as the driver of hyperglycemia (Kraegen et al., 1991; Turner et al., 2013).

### Limitation

The identities of nutrient storages were not explicitly studied in this work, but determined based on literature knowledge. For example, triglycerides and proteins are storage for NEFAs and amino acids, respectively. However, while it is commonly assumed that glycogen is the storage for glucose, we recognize literature evidence pointing to other forms of tissue storages for glucose, such as proteins (Shipley et al., 1967, 1970). Furthermore, as glucose is a substrate for the glycerol backbone of tissue TAG, there is likely substantial flux from glucose to this form of nutrient storage. A thorough investigation of the storages with new experimental designs would be a critical future objective. In addition, our list of circulating nutrients did not include circulating TAG; despite its larger pool size than NEFA, TAG has a small circulatory turnover flux (Gormsen et al., 2006; Lambert et al., 2014; Sarac et al., 2012), and accounts for a minor fraction of total lipid oxidation (Diraison and Beylot, 1998). As VLDL-TAG is mostly derived from circulating NEFA and was labeled by ^13^C-FA tracer during the time frame of our *f*_*ox*_ measurement (Parks and Hellerstein, 2006; Parks et al., 1999), its oxidation and storage fluxes were mostly accounted for in the NEFA fluxes, especially for lean mice. Future work is needed to explicitly delineate the NEFA and TAG fluxes. Finally, our work focused on establishing the flux framework and for this purpose only studied fasted male mice. Application of this framework to female animals and to other physiological conditions such as feeding is of great importance.

## ACKNOWLEDGEMENTS

The authors thank all current members and alumni of the Hui lab for their assistance and discussion in the research. This research was supported by the Paul G. Allen Family Foundation 0034665 (S.H.), the NIH R00DK117066 (S.H.), and the Sabri Ülker Center for Metabolic Research. Some figures were created with BioRender.com.

## AUTHOR CONTRIBUTIONS

B.Y. designed and conducted the experiment, analyzed data, and wrote the manuscript. O.T. and W.D. performed the hyperinsulinemia clamp, ITT and GTT. Y.Y.K. performed the lipolysis enzymatic activity assay. K.I. supervised and assisted with the metabolic cage measurement. G.H. co-supervised the metabolic cage measurement and proposed critical revision to the manuscript. S.H. conceived and supervised the project, secured funding, and wrote the manuscript. All authors reviewed and contributed feedback to the manuscript.

## STAR METHODS

### KEY RESOURCES TABLE

| REAGENT or RESOURCE | SOURCE | IDENTIFIER |
| --- | --- | --- |
| **Chemicals, Peptides, and Recombinant Proteins** |  |  |
| D-glucose ($^{14}\text{C}_6$) | PerkinElmer | NEC042V250UC |
| D-glucose ($^{13}\text{C}_6$, 99%) | Cambridge Isotope Laboratories | CLM-1396 |
| sodium L-lactate ($^{13}\text{C}_3$, 98%) 20% w/w in $\text{H}_2\text{O}$ | Cambridge Isotope Laboratories | CLM-1579 |
| L-alanine ($^{13}\text{C}_3$, 99%) | Cambridge Isotope Laboratories | CLM-2184-H |
| L-glutamine ($^{13}\text{C}_5$, 99%) | Cambridge Isotope Laboratories | CLM-1822-H |
| glycerol ($^{13}\text{C}_3$, 99%) | Cambridge Isotope Laboratories | CLM-1510 |
| sodium D-3-hydroxybutyrate ($^{13}\text{C}_4$, 99%) | Cambridge Isotope Laboratories | CLM-3853 |
| sodium acetate ($^{13}\text{C}_2$, 99%) | Cambridge Isotope Laboratories | CLM-440 |
| sodium palmitate ($^{13}\text{C}_{16}$, 98%) | Cambridge Isotope Laboratories | CLM-6059 |
| oleic acid, potassium salt ($^{13}\text{C}_{18}$, 98%) | Cambridge Isotope Laboratories | CLM-8856 |
| linoleic acid, potassium salt ($^{13}\text{C}_{18}$, 98%) | Cambridge Isotope Laboratories | CLM-8835 |
| L-valine ($^{13}\text{C}_5$, 99%) | Cambridge Isotope Laboratories | CLM-2249-H |
| bovine serum albumin, fatty acid free, $\geq 96\%$ | Sigma-Aldrich | A6003 |
| glycerokinase | Sigma-Aldrich | G6278 |
| Ultra Sensitive Mouse Insulin ELISA Kit | Crystal Chem | 90080 |
| **Experimental Models: Organisms/Strains** |  |  |
| Mouse: C57BL/6J | Jackson laboratory | 000664 |
| Mouse: ob/ob <HO> | Jackson laboratory | 000632 |
| **Software and Algorithms** |  |  |
| El-MAVEN software | Elucidata https://docs.polly.elucidata.io/index.html |  |
| RStudio | Posit https://posit.co/download/rstudio-desktop/ |  |
| AccuCor | Github: https://github.com/XiaoyangSu/AccuCor |  |
| Biorender | Biorender https://www.biorender.com/ |  |
| **Others** |  |  |
| chow diet | PicoLab Mouse Diet 20 | 5058 |
| high fat diet | Research Diets | D12492i |
| XBridge BEH Amide XP column | Waters | 176002889 |
| Acquity UPLC BEH C18 column | Waters | 186003719 |
| catheter for mouse jugular vein | Instech Laboratories | C20PUMJV1301 |
| catheter for mouse carotid artery | Instech Laboratories | C10PU-MCA1459 |
| vascular access button, 1 channel | Instech Laboratories | VABM1B/25 |
| vascular access button, 2 channels | Instech Laboratories | VABM2B/25R25 |
| tether for 1 channel button | Instech Laboratories | VABM1T/25 |
| tether for 2 channel button | Instech Laboratories | VABM2T/25 |
| Buprenorphine base ER 0.5 mg/mL | Wedgewood | Rx # 7270862 |
| Swivel | Instech Laboratories | 375/25PS |
| counter balanced arm | Instech Laboratories | SMCLA |
| Blood collection Microvette | Starstedt Inc | 16.440.100 |
| Heparin sodium 5000 U/mL | McKesson Corp. | - |

### RESOURCE AVAILABILITY

#### Lead Contact

Additional information and requests for resources should be directed to the Lead Contact, Sheng Hui.

#### Materials AVAILABILITY

No new reagents were generated in this study.

#### Data and Code Availability

Raw data and R scripts used for flux analysis is available on GitHub (https://github.com/yuanboFaith/flux-model-energy-metabolism).

### EXPERIMENTAL MODEL AND SUBJECT DETAILS

All mouse work followed protocols approved by the Institutional Animal Care and Use Committee of Harvard Medical Area. The animals were housed on a normal light cycle (7 AM to 7 PM), and were 12-24 weeks old when used in experiment. The wild type C57BL/6J and ob/ob mice were fed with standard chow (PicoLab Mouse Diet 5058). For diet-induced obese mice, C57BL/6J mice were fed with high fat diet (Research Diets D12492i, 60%kcal from fat) starting from the 3 weeks of age for 3-4 months before experiment. For *in vivo* tracer infusion, a jugular vein catheter (JVC) was inserted into the right jugular vein, under aseptic conditions and with anesthesia using isoflurane. The JVC was tunneled subcutaneously to the back of the interscapular area, and connected to a single-channel vascular access button (VAB). For arterial blood collection, in addition to jugular catheterization, a carotid artery catheter (CAC) was implanted into the left carotid artery. Both the JVC and CAC were subcutaneously tunneled to the interscapular region, and connected to a double-channel VAB. The catheter patency was maintained by sealing with lock solution composed of 50% glycerol in saline, with 200 U heparin/mL. After surgery, 50 µL of 0.5 mg/mL analgesic buprenorphine (extended release) per 25 g body weight was subcutaneously injected to the right flank. The animal was allowed to recover from surgery for at least five days before use in experiment.

### METHOD DETAILS

#### ^13^CO_2_ Measurement

^13^C-labeled tracer compounds were prepared in sterile saline. Fatty acids were complexed with bovine serum albumin in a molar ratio of 6:1. For ^13^CO_2_ measurements after tracer bolus injection, the mouse was transferred to a new cage without food at 9:00 AM. After 4 hours of fasting, the animal was gently warmed by a heat lamp to dilate the tail vein. Under light isoflurane anesthesia, the mouse was administered with 200 µL ^13^C-tracer solution via the tail vein. The animal was then immediately transferred to a cage in the metabolic cage system, which was equipped with a stable isotope gas analyzer (Promethion, Sable Systems International, NV). Expired ^13^CO_2_ and metabolic cage parameters including air flow rate, oxygen, and CO_2_ were monitored for another 6 hours without food access.

^13^CO_2_ measurement with tracer infusion required catharized mice and followed the same protocol described in the tracer infusion section below, except that the infusion lasted for 3-4 hours, with a reduced tracer concentration. The tracer concentration and infusion rates can be found in **Table S1**. ^13^CO_2_ data and metabolic cage parameters including air flow rate, oxygen, and CO_2_ were continuously collected for the entire period of tracer infusion.

#### Infusion of Isotope Tracers

On the day of tracer infusion, the mouse was transferred to a new cage without food at 9 AM, and fasted for 4 hours before infusion. Right before the start of infusion, the lock solution was drawn and removed from the catheter, and the catheter was flushed with 50 µL saline. The VAB was then connected to the infusion tether, which was in turn connected with a swivel on a counter-balance arm. This allowed the mouse to move freely around in the cage. The infusion of U-^13^C-tracer at a small non-perturbing dose started at 2 PM, and lasted for 30 min for fatty acids and 2.5-3 h for all other tracers. The tracer concentration and infusion rates can be found in **Table S1**. At the end of infusion, 40 µL blood was collected by tail snip. For infusion of ^13^C-lactate, alanine, and glutamine, given their significant labeling difference in the arterial and venous blood (**Table S2**)(Lee et al., 2023), 40 µL arterial blood was sampled from the carotid artery via the pre-implanted catheter, and used for analysis. The blood was collected into a Microvette tube with clotting activator, and was kept on ice during collection. After infusion, the catheter was flushed with 50 µL saline, and then sealed with 10 ∼ 20 µL lock solution. The blood was centrifuged at 5,000 ×g for 10 min. 4 µL serum was aliquoted in separate tubes, and stored at -80°C.

#### Hyperinsulinemia Clamp

BL/6J mouse (26.5 ± 1.8 g) was transferred to a new cage without food at 9 AM. After fasting for 4 hours, blood was collected via tail snip for basal glycemia and insulin measurement. A mixture of U-^13^C-glucose (150 mM), unlabeled glucose (2361 mM) and insulin (66.6 mU/mL) was infused at 3.3 µL/min/mouse for 2.5 h, and blood was collected for serum glucose labeling and insulin measurement. Then U-^13^C palmitate (10.68 mM) was infused at 6 µL/min/animal for another 30 min (delivered by a second pump), simultaneously with the glucose and insulin infusion. Blood was collected at the end for serum glucose and palmitate labeling and insulin measurement. Insulin was measured by ELISA Kit following the vendor’s protocol.

#### Serum Sample Preparation for LC-MS Analysis

4 µL serum was mixed with 60 µL -20°C methanol: acetonitrile: water at 40: 40: 20. The sample was vigorously vortexed, and centrifuged at 4°C at 13,000 g for 10 min. 30 µL supernatant was transferred to the HPLC vial from which 5 µL was injected to the LC-MS.

Serum glycerol was first enzymatically derivatized into glycerol-3-phosphate before LC-MS analysis. 4 µL serum was mixed with 90 µL freshly made enzyme solution containing 2 U/mL glycerol kinase, 25 mM Tris-HCl (pH 8.0), 50 mM sodium chloride, 10 mM magnesium chloride, and 1.5 mM ATP, and incubated at room temperature for 15 min. The reaction was stopped by the addition of 400 µL methanol. The sample was vortexed, and centrifuged at 16,000 g for 10 min. The supernatant was transferred to a new tube, and dried using SpeedVac at 45 °C for 90 min. 60 µL of methanol: acetonitrile: water at 40: 40: 20 was added to the residue, vortexed, and centrifuged at 16,000 g for 10 min. 30 µL supernatant was transferred to the HPLC vial, from which 5 µL was injected to the LC-MS.

#### LC-MS Analysis

Chromatographic separation was achieved using XBridge BEH Amide XP Column (2.5 µm, 2.1 mm × 150 mm) with guard column (2.5 µm, 2.1 mm × 5 mm) (Waters, Milford, MA). The mobile phase A was water: acetonitrile 95:5, and mobile phase B was water: acetonitrile 20:80, with both phases containing 10 mM ammonium acetate and 10 mM ammonium hydroxide. The elution linear gradient was: 0 ∼ 3 min, 100% B; 3.2 ∼ 6.2 min, 90% B; 6.5. ∼ 10.5 min, 80% B; 10.7 ∼ 13.5 min, 70% B; 13.7 ∼ 16 min, 45% B; and 16.5 ∼ 22 min, 100% B, with flow rate of 0.3 mL/min. The autosampler was at 4°C. The injection volume was 5 µL. Needle wash was applied between samples using methanol: acetonitrile: water at 40: 40: 20. The mass spectrometer used was Q Exactive HF (Thermo Fisher Scientific, San Jose, CA), and scanned from 70 to 1000 *m/z* with switching polarity. The resolution was 120,000. Metabolites were identified based on accurate mass and retention time using an in-house library. ^13^C-Natural abundance correction was performed in R using package AccuCor (Su et al., 2017).

#### ^14^C-glucose administration and carcass analysis

The mouse was fasted from 9 AM to 12 PM, and then 12 nCi ^14^C_6_-glucose in 200 µL saline was administered via the tail vein. After 6 h, the animal was sacrificed, and the hair was shaved off. The carcass was transferred into an Erlenmeyer flask containing 70 mL 1.7 M KOH, and microwaved to simmer. The flask was incubated at 90 C° for 3 h for complete carcass digestion. After having cooled down to room temperature, the total liquid volume was measured, and 2 mL 15% antifoam B emulsion (Sigma A5757) and 1 mL Tween 80 (Sigma P4780) was added to the digest. Under magnetic stirring, 0.4 mL of the digest was sampled and transferred to a scintillation vial. A mix of 150 µL 2 M acetic acid, 200 µL 50% H_2_O_2_, and 50 µL 15% antifoam B was added to the vial, followed by incubation at 70C° for 20 min to decolorize the mixture. The vial was then added with 200 µL of 12 M acetic acid and 5 mL Ecolume scintillation liquid (MP Biomedicals, 0188247001) and vigorously vortexed. ^14^C radioactivity in the vial was measured using a HIDEX 300 SL liquid scintillation counter (Turku, Finland).

### QUANTIFICATION AND STATISTICAL ANALYSIS

#### Calculation of the Fraction of Oxidation (*f*_*ox*_) of a Circulating Nutrient

The *f*_*ox*_ of a circulating nutrient was calculated as the fraction of ^13^C atoms in the exhaled CO_2_ among the total ^13^C atoms in the administered tracer. For a mouse in a metabolic cage system administered with a ^13^C tracer, the production flux of tracer-derived ^13^CO_2_ from the animal *J* (µmol/min) was calculated as

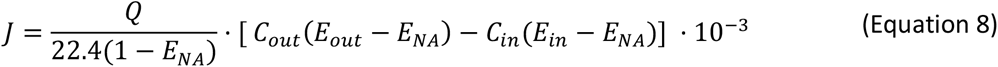

where *Q* is the air flow rate through the cage, *C*_*in*_ and *C*_*out*_ are the CO_2_ concentration measured in the air inflow and outflow respectively, *E*_*in*_ and *E*_*out*_ are the ^13^CO_2_ enrichment measured in the air inflow and outflow, respectively, and *E*_*NA*_ is the ^13^C natural abundance of the unlabeled nutrients in the mouse body. *E*_*NA*_ was calculated using Eq. (8) from measured CO_2_ concentration and enrichment data for a non-infused control mouse group and a production flux of tracer-derived ^13^CO_2_ as zero (*J* = 0). For an animal under the infusion at the rate of *I* (µmol ^13^C-atoms/min), the *f*_*ox*_ was then calculated as

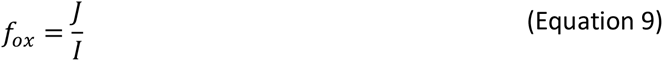

For animal receiving the tracer as a single bolus at dosage of *B* (µmol ^13^C-atoms), the fraction of oxidation *f*_*ox*_ was calculated as

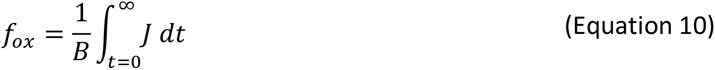

with integration from the time of injection to the end of measurement when CO_2_ enrichment reached basal level. Details of the derivations are in **Suppl. Note 1**.

#### Calculation of Fluxes to Oxidation (*J*_*OX*_) and Storages (*J*_*S*_) from a Circulating Nutrient

The *J*_*ox*_ and *J*_*S*_ were calculated using Eqs. (1) and (2) in the main text, respectively. The labeling *L* used here and throughout the text was the carbon-atom labeling fraction of the nutrient. It is calculated as

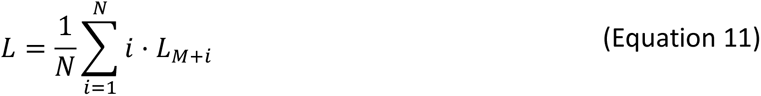

where *N* is the number of carbon atoms in a molecule, and *L*_*M*+*i*_ is the fraction of isotopologue containing *i* number of ^13^C atoms among all isotopologues of the molecule. Details of the derivations are in **Suppl. Note 2**.

#### Calculation of the *<u>Direct</u>* Fluxes to Oxidation (*O*) and Storage (*S*) from Circulating Nutrients

The <u>direct</u> oxidation fluxes *O*_*j*_ of *N* circulating nutrients were calculated with Eq. (5) in the main text. The direct storing fluxes *S*_*j*_ of the circulating nutrients were calculated with

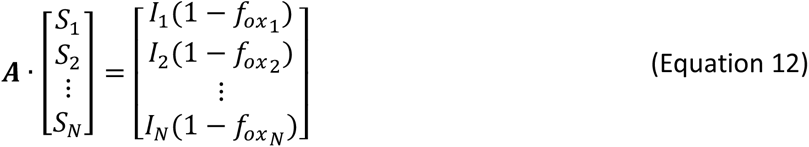

where *I*_*j*_ and 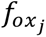 are the infusion rate and the fraction of oxidation of nutrient *j*, respectively.

The matrix ***A*** is defined as

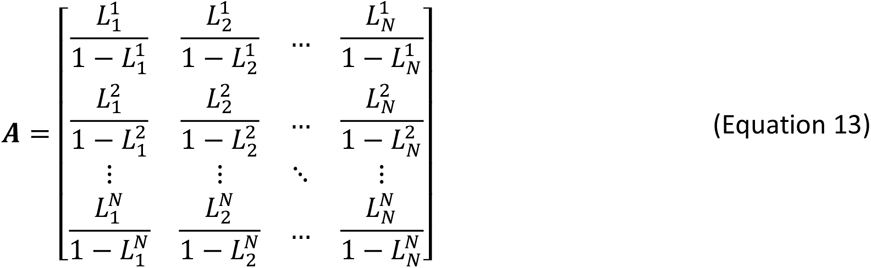

where 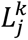 is the labeling of nutrient *j* under the infusion of ^13^C-labeled nutrient *k*. The unknowns *O*_*j*_ and *S*_*j*_ were solved with the method of least squares, subjected to *O*_*j*_ ≥ 0 and *S*_*j*_ ≥ 0, using R package LimSolve, and function lsei (least squares with equalities and inequalities) (Soetaert, 2009). The error was approximated with Monte Carlo simulation. For Eqs. (5) and (12), respectively, we ran the matrix optimization 100 times. In each optimization, for each 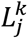 and 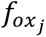 value, a random value was drawn from a normal distribution, with mean and standard deviation (SD) being the mean and standard error of the mean (SEM) of these measured quantities. The SD of the 100 simulated ***O*** and ***S*** vectors were taken as the associated SEM, respectively. More details of the calculation can be found in **Suppl. Note 3**.

#### Calculation of the Storage Releasing Flux (*F*_*s*_) to a Circulating Nutrient, and Conversion Fluxes (*F*_*j*_) between Circulating Nutrients

To calculate the storage-releasing flux *F*_*S*_ and the converting fluxes *F*_*j*_ from *n* different circulating nutrients to a target nutrient (TN) of interest, ^13^C-labeled TN and the *n* source nutrients were infused individually to the animals. Each infusion generates an Eq. (7), with *L*_*TN*_ in replace of *L*_*glu*_.

Integrating all *n* + 1 equations gives the linear system

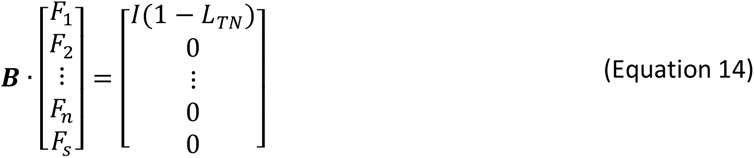

with

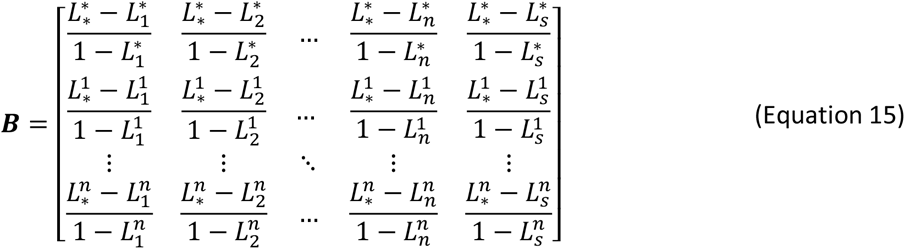

where 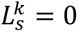. Asterisks are used in replace of *TN* for notation clarity. The matrix optimization and error estimation were performed as above. More details of the calculation can be found in **Suppl. Note 4**.

#### Estimation of Energy Wasted in Futile Cycles

The ATP flux that is wasted in the major futile cycles was estimated using the following parameters: 4 ATP molecules wasted per cycling of one glucose molecule in the Cori cycle, 9-11 ATP molecules used in the resynthesis of each TAG molecule in the TAG-NEFA cycling, and 4 ATP molecules used for synthesis of each peptide bond (Bender, 2012). The total flux of ATP generation was calculated based on the direct carbon oxidation flux of circulating nutrients, where the ATP yield is 5.3 per carbohydrate carbon, 6.5 per fatty acid carbon, 5.5 per ketone body carbon, and 5 per amino acid carbon (Bender, 2012). More details of the calculation are in **Suppl. Note 6**.

