## Supplemental Figures for "An Organism-Level Quantitative Flux Model of Energy Metabolism in Mice"

### Supplementary Figures

Bo Yuan [1], Will Doxsey [1], Özlem Tok [1], Young Yon Kwon [1],
Karen E. Inouye [1,2], Gökhan S. Hotamışlıgil [1,2,3], Sheng Hui [1,2,4,*]

[1] Department of Molecular Metabolism, Harvard T. H. Chan School of Public Health, Boston, Massachusetts, USA.

[2] Sabri Ülker Center for Metabolic Research, Department of Molecular Metabolism, Harvard T.H. Chan School of Public Health, Boston, Massachusetts, USA.

[3] Broad Institute of Harvard and MIT, Cambridge, Massachusetts, USA.

[4] Lead Contact

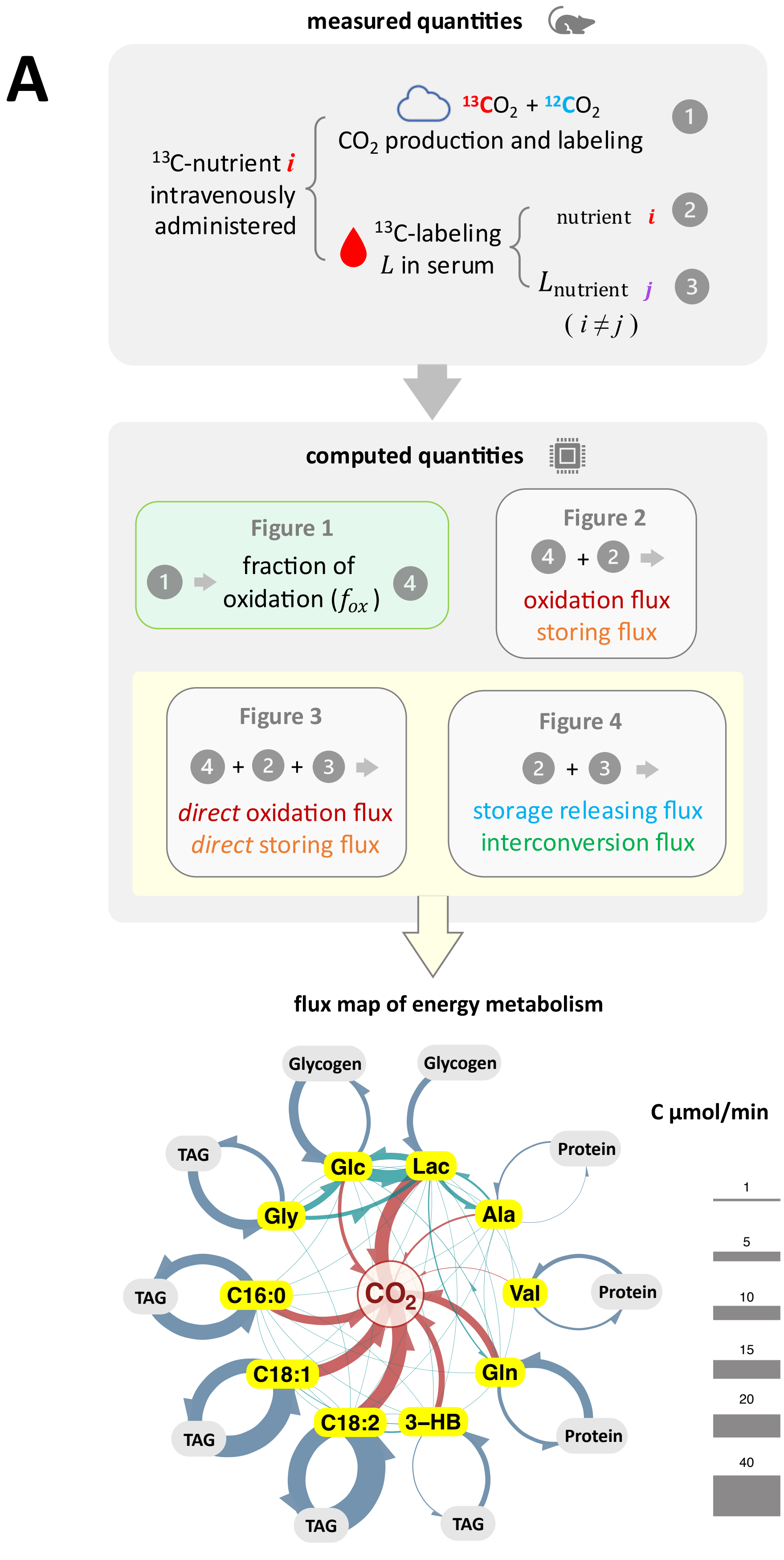


Figure S1 (continued)

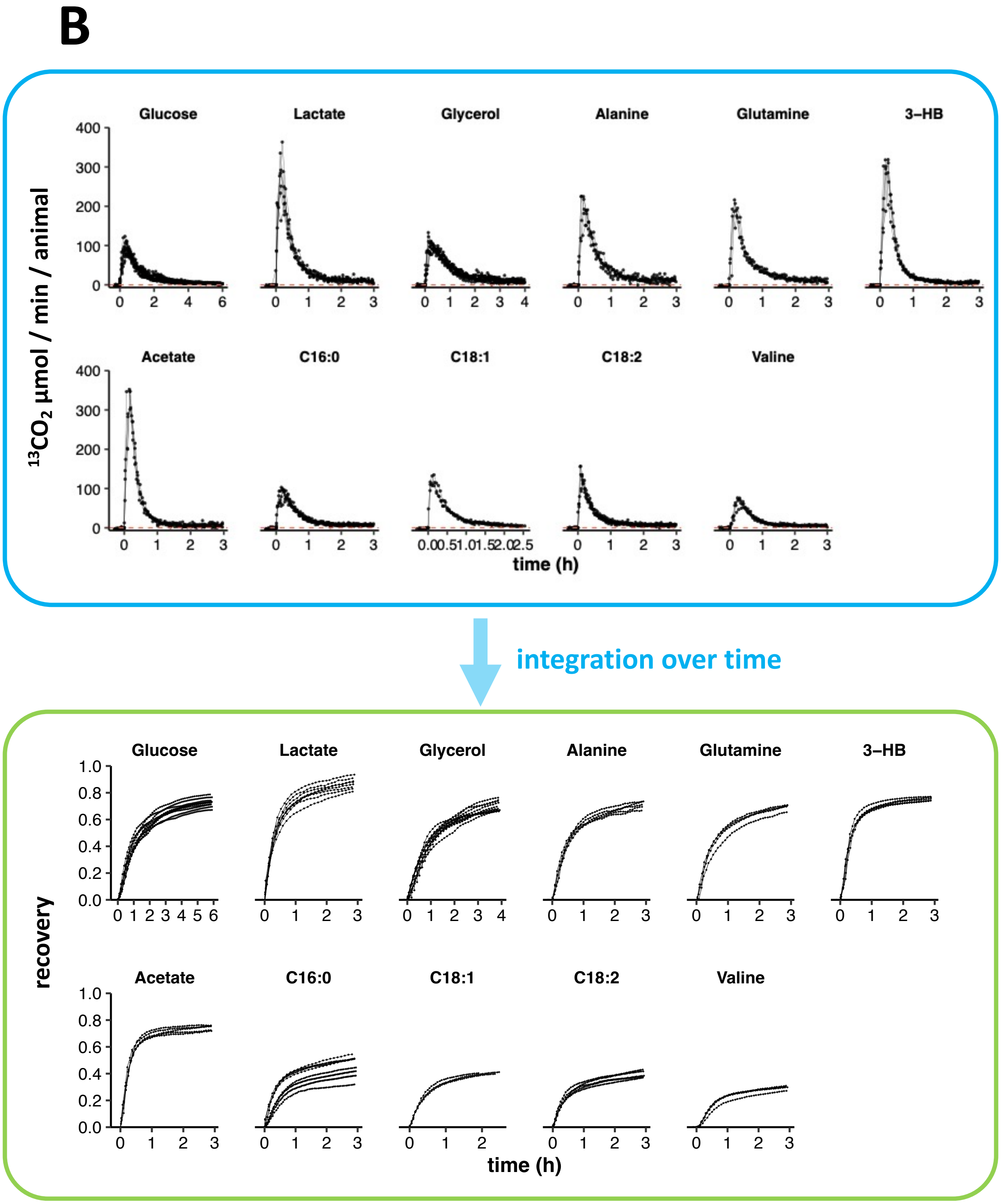


Figure S1 (continued)

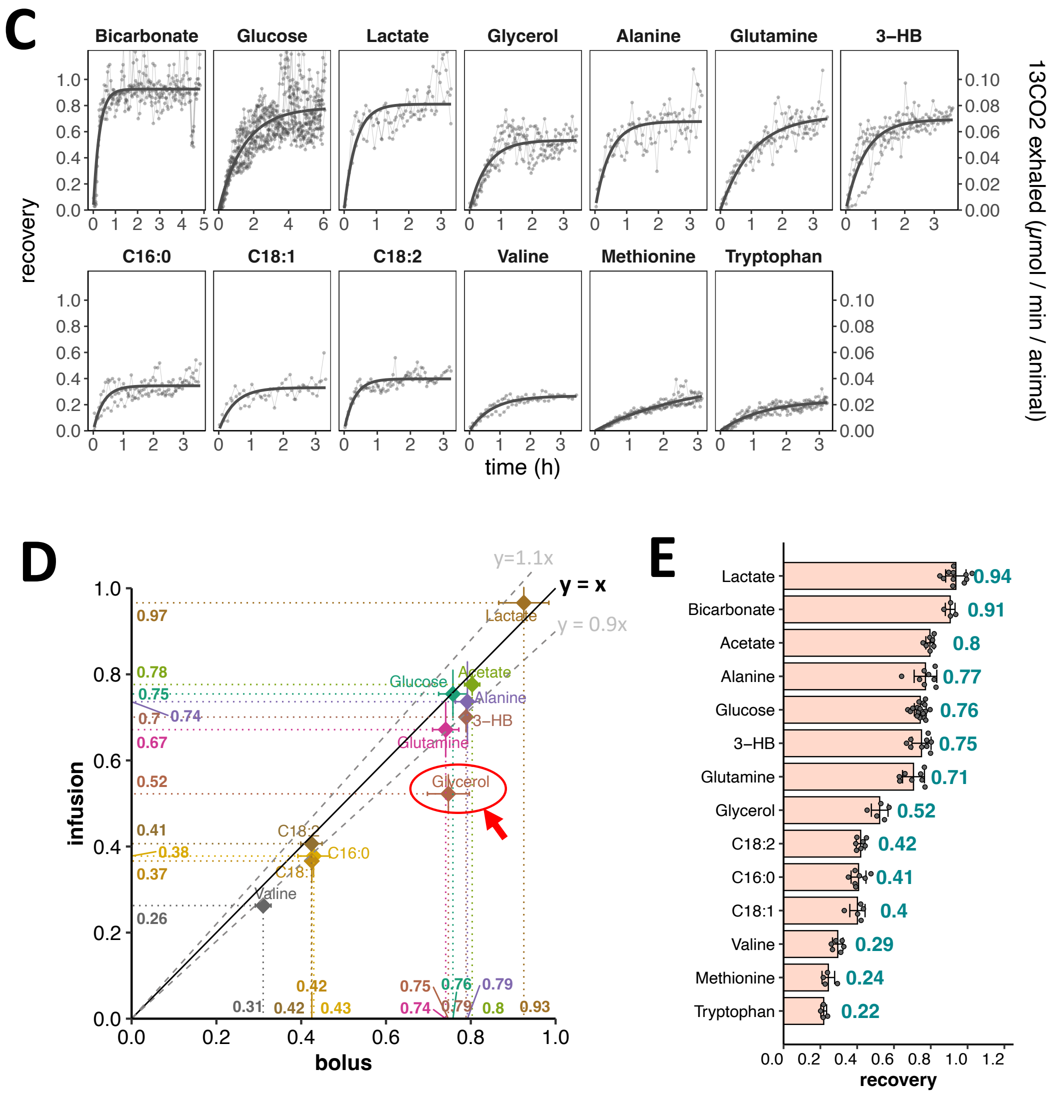


**Figure S1. $^{13}CO_2$ Tracing Unveils Characteristic Oxidation and Storing Partition of Circulating Nutrients. Related to Figure 1.**

**(A)** Workflow to measure and calculate metabolic fluxes of circulating nutrients.

**(B)** Instant $^{13}CO_2$ production rate (top) after $^{13}C$ bolus injection, and the accumulated recovery of $^{13}C$ in exhalation relative to the administered dosage (bottom). Data is normalized to administration of 10 μmol $^{13}C$-atoms. The data shown is before the correction of bicarbonate recovery. Each connected line is a biological replicate, n = 3 to 10 for each nutrient.

**(C)** Recovery of $^{13}C$ during $^{13}C$-nutrient infusion with non-perturbative dose in conscious non-restrained mice. Data is normalized to infusion rate of 0.1 μmol $^{13}C$ / min / animal. The regression curve is fitted with formula $y = a \cdot (1 - e^{-b \cdot t})$. Each connected line is a biological replicate, n = 2 to 10 for individual nutrient.

**(D)** Comparison of the $^{13}C$ recovery by two $^{13}C$-administration methods, bolus injection vs. continuous infusion.

**(E)** $^{13}C$ recovery of circulating nutrients. The data is pooled from data by bolus administration (Panel B) and infusion (Panel C). Only infusion data is used for glycerol.

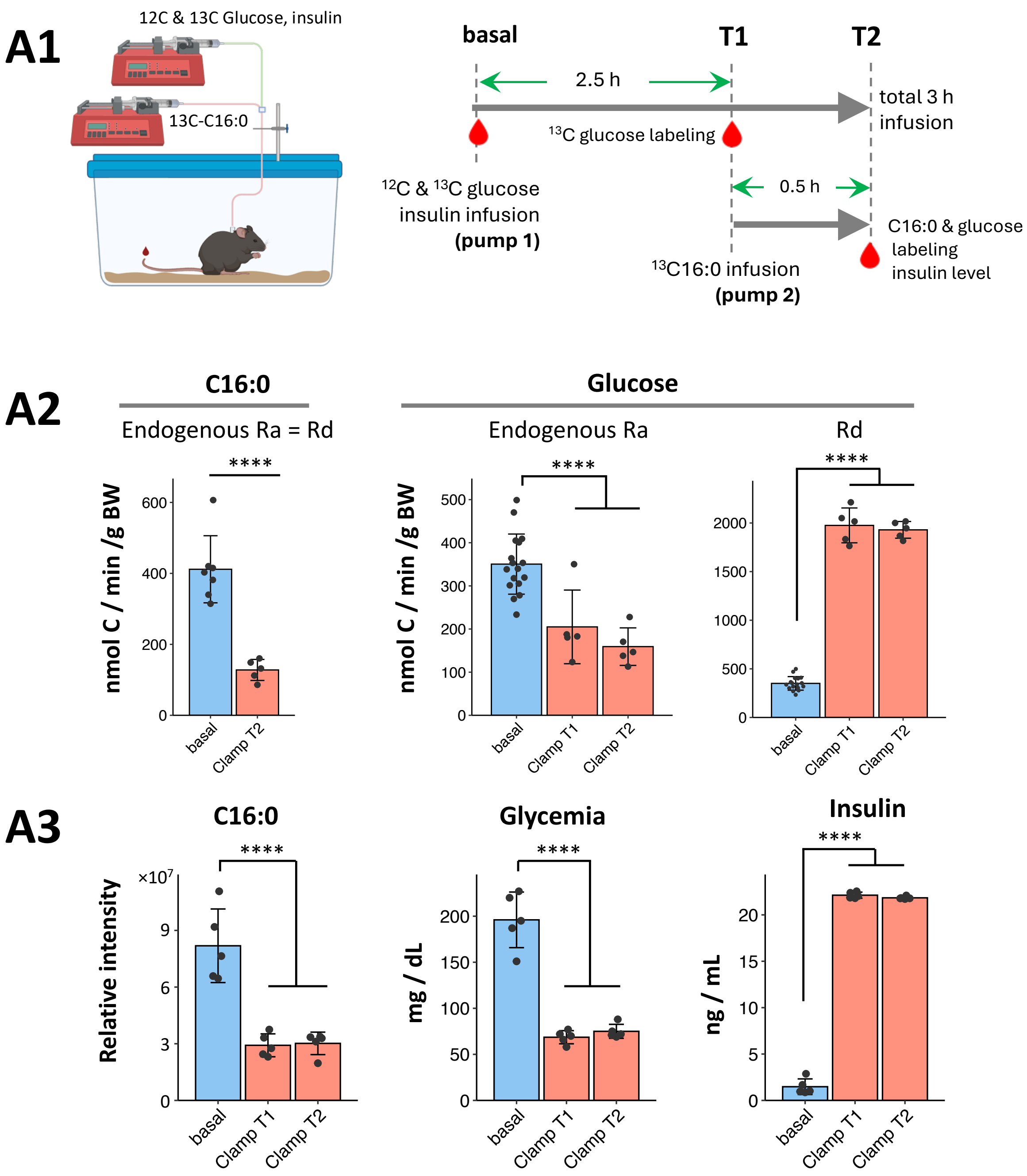


Figure S2 (continued)

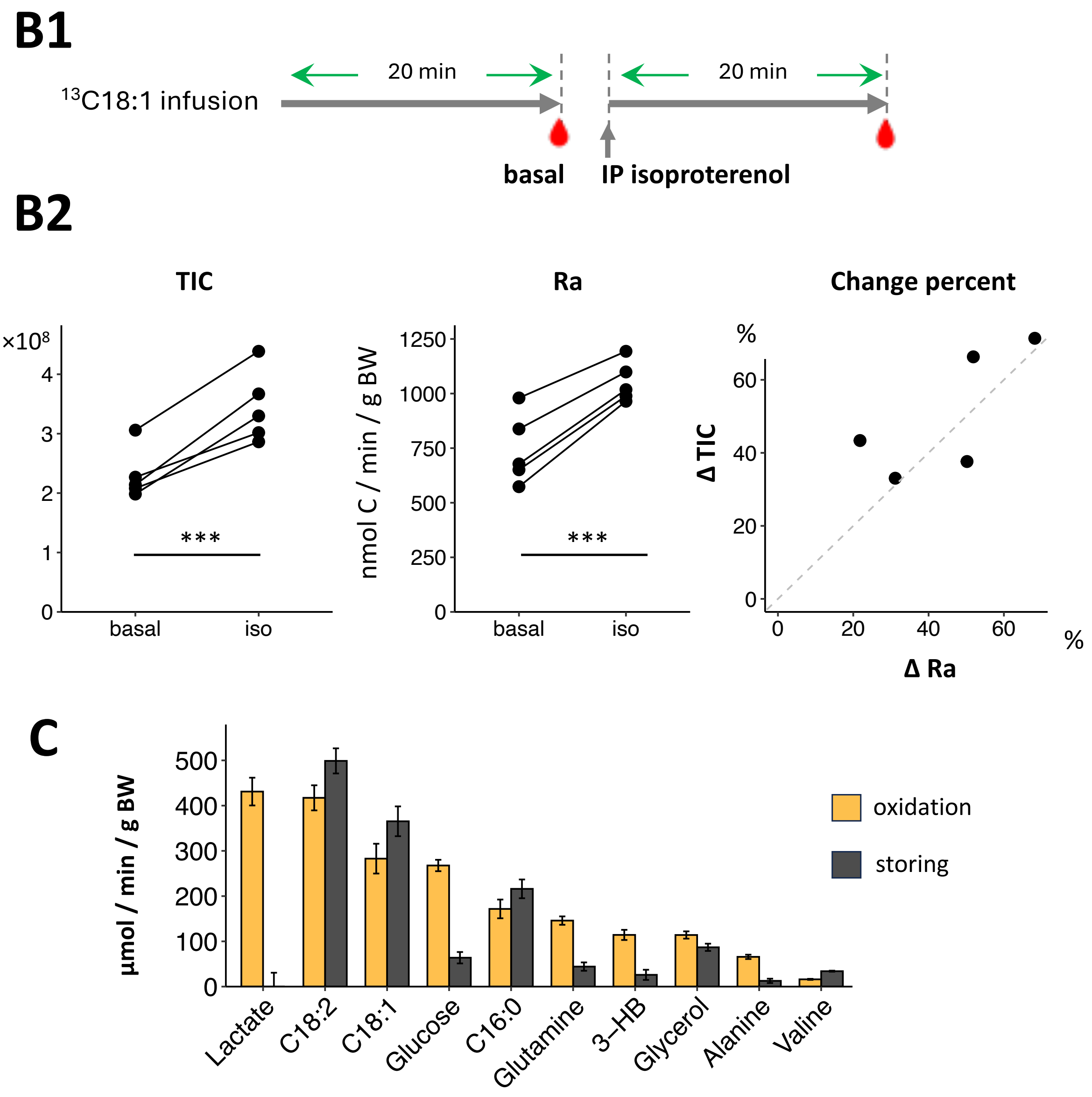


**Figure S2. Circulating Nutrients' Endogenous Production, Consumption, Oxidation, And Storage Flux at Basal and Varied Physiological Conditions. Related to Figure 2.**

**(A)** Hyperinsulinemia suppresses lipolysis and endogenous glucose production, and increases glucose disposal. *(A1)* Scheme of experimental design of hyperinsulinemia clamp. *(A2)* Rate of appearance (Ra) and disposal (Rd) of palmitate (C16:0) and glucose at basal and hyperinsulinemia condition. The basal flux was measured in separate experiments. *(A3)* Blood palmitate, glucose and insulin level at basal and hyperinsulinemia condition.

**(B).** Isoproterenol increases lipolysis. *(B1)* Scheme of experimental design. *(B2)* Serum oleate relative abundance (total ion counts, TIC, left plot), rate of appearance (Ra, middle plot), and correlation between percent of change between TIC and Ra (right plot). For panels (A-B), statistics followed paired t-test.

**(C)** Overall oxidation and storage fluxes of major circulating nutrients at basal fasting condition. mean ± sem. Flux is normalized by body weight. Flux at whole-body level refers to Figure 2C.

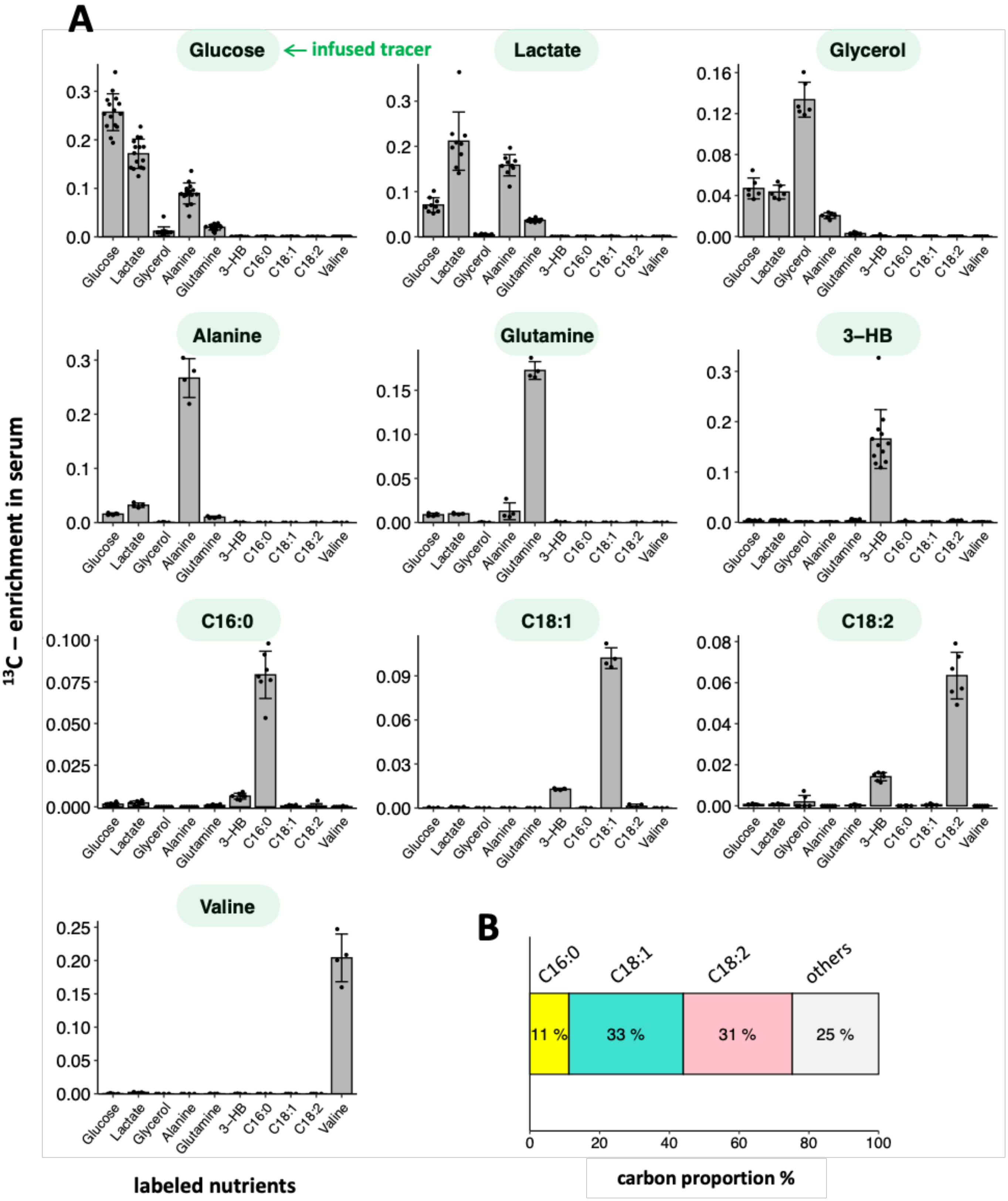


Figure S3 (continued)

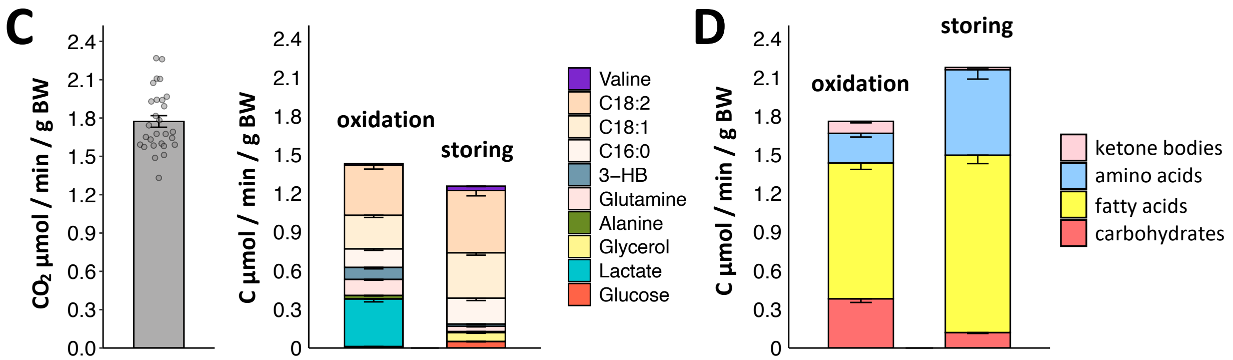


**Figure S3. Supporting Data used for Quantifying the *Direct* Oxidation and Storing Fluxes of Circulating Nutrients. Related to Figure 3.**

**(A)** The $^{13}$C labeling of serum nutrients at the isotopic steady state of infusion of different $^{13}$C-nutrients. n = 4 to 15 for individual nutrient, noted in the scatterplot.

**(B)** Carbon atom distribution in non-esterified fatty acids in serum. The analytes were determined using LC-MS (see Methods section). n = 6.

**(C)** Total CO2 production (left), in comparison with the direct oxidation and storing fluxes of the major circulating nutrients covered in this study (right). Related to Figure 3C.

**(D)** Direct oxidation and storing fluxes of all major circulating nutrients, including carbohydrates (glucose, lactate, and glycerol), fatty acids, protein-originated amino acids, and ketone bodies. Related to Figure 3D.

All data is presented in mean ± SEM. Fluxes in panel (C-D) are in unit of µmol / min / g body weight (BW), and fluxes per animal refers to Figures 3 C-D.

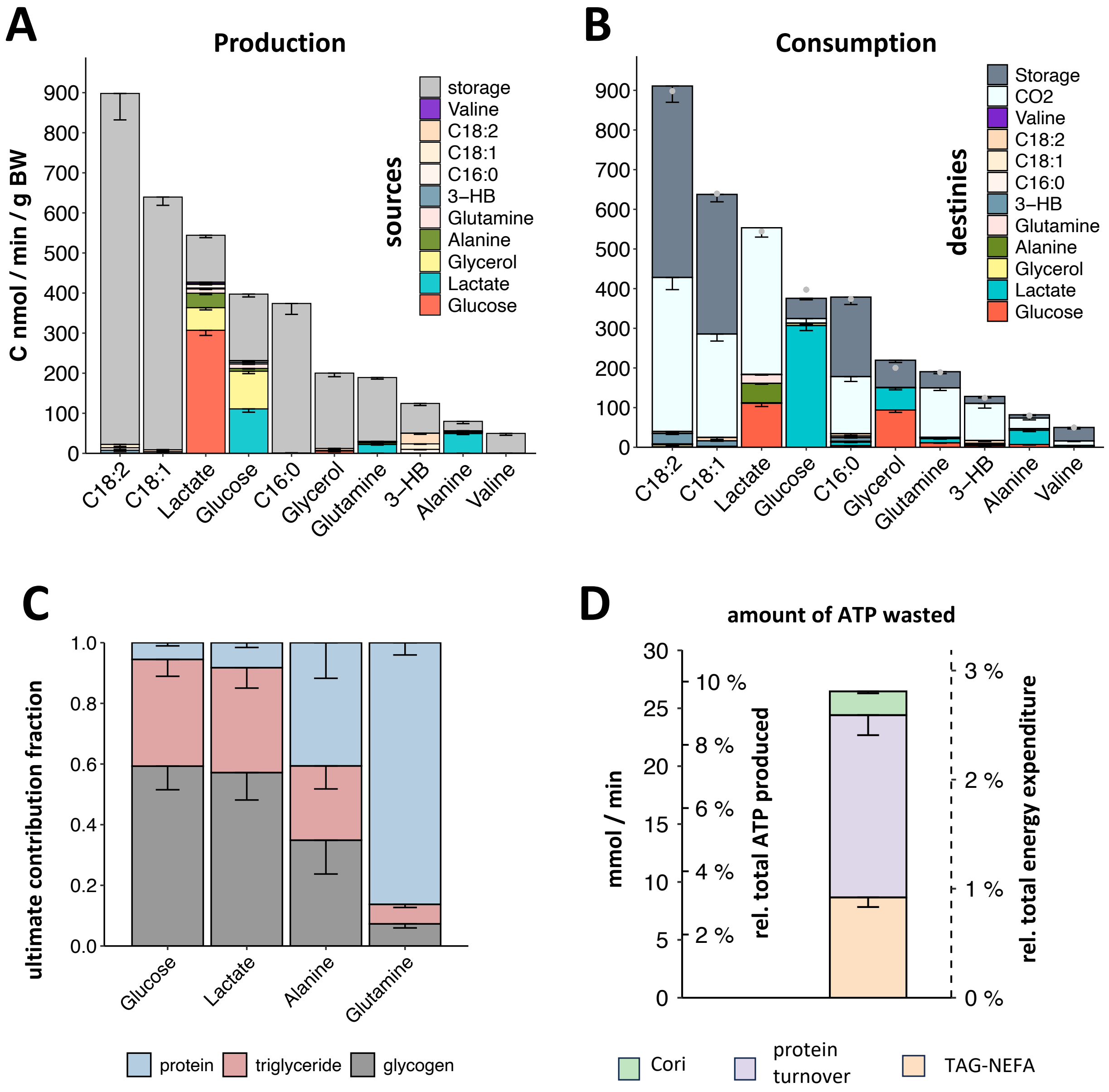


**Figure S4. Quantifying Storage Releasing Flux and Interconverting Fluxes for Circulating Nutrients, and Integrating them with Oxidation and Storing Fluxes into a Holistic Flux Model of Circulating Nutrients. Related with Figure 4.**

**(A)** Production fluxes of circulating nutrients from storage and other circulating nutrients. Flux is normalized by body mass. Fluxes per animal is presented in Figure 4B.

**(B)** Sink fluxes of circulating nutrients, fated to oxidation, storage, and other circulating nutrients. Fluxes are normalized by body mass. Fluxes per animal is presented in Figure 4C.

**(C)** The ultimate contribution fraction from storages to circulating nutrients. Details of the calculation are in **Suppl. Note 5**.

**(D)** Amount of ATP wasted by the top 3 futile cycles.

Data is presented in mean ± SEM.

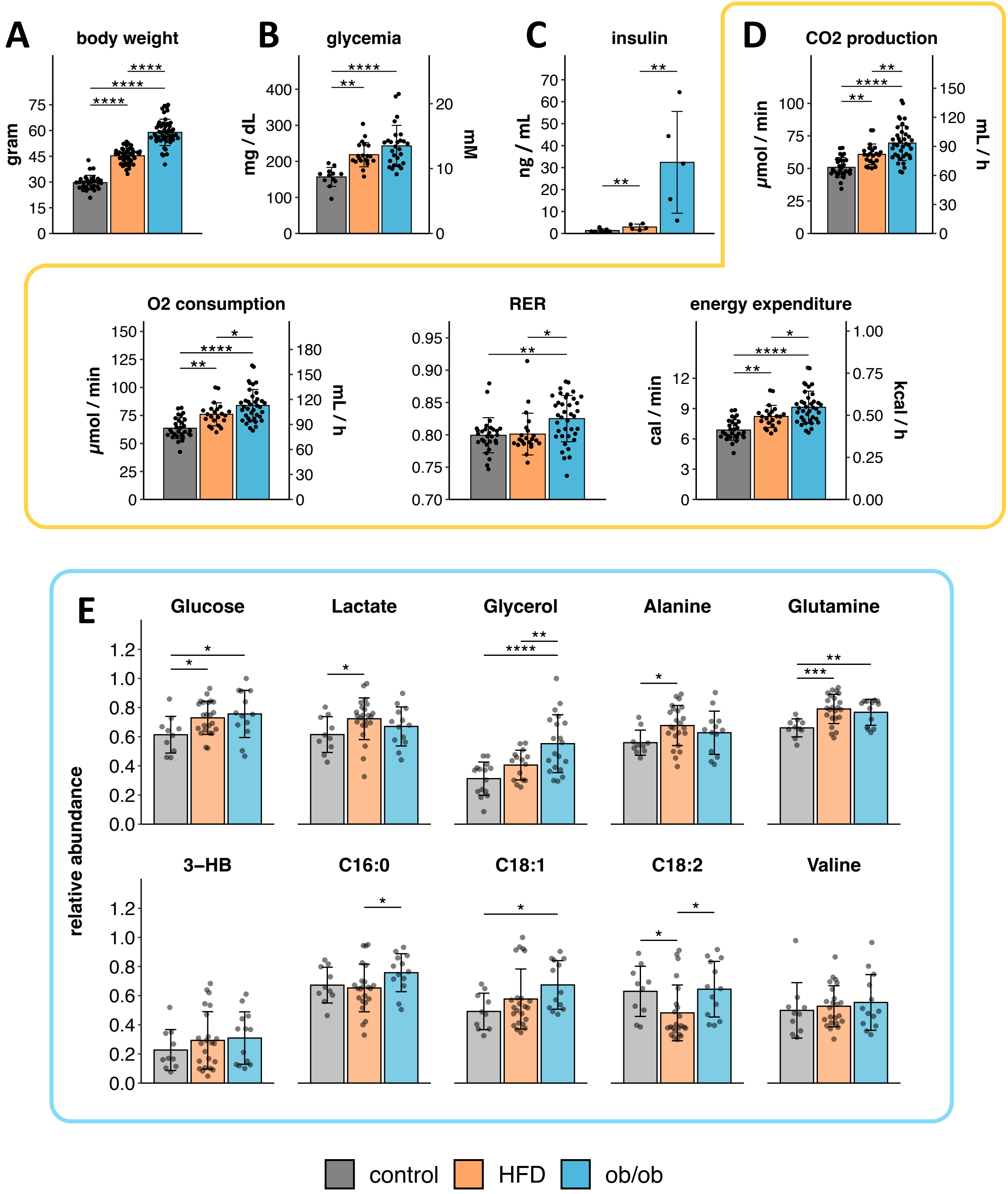


**Figure S5. Basic Phenotypes of the Control and Obese Mice. Related to Figure 5.**

**(A)** The body weight of the animals.

**(B)** Blood glucose level measured by glucometer.

**(C)** Serum insulin level.

**(D)** Metabolic cage measurement, including $CO_2$ production, $O_2$ consumption, respiratory exchange ratio (RER), and energy expenditure (EE). Data shown is from $^{13}C$-infusion experiment for $^{13}CO_2$ measurement. In this panel, multiple measurements separated by a week on the same mouse are counted as biological replicates. Note that for $CO_2$ production,

$O_2$ consumption and EE, when the body weight is adjusted with analysis of covariance (ANCOVA)(Mina et al., 2018), the difference in phenotypes is no longer significant. This suggests that the apparent differences in whole-body energy expenditure is likely a result of differences in body size, instead of intrinsic metabolic differences.

**(E)** Relative abundance of circulating nutrients in serum measured by LC-MS. The data is from the $^{13}C$-infusion experiment for serum labeling. On a nutrient-wise basis and separately for each MS sequence, the peak intensity is first normalized as fraction of the largest peak intensity, and then data of each mouse is averaged across different infusion experiments.

All data is presented as mean ± SD. *$p < 0.05$, **$p < 0.01$, ***$p<0.001$, ****$p<0.0001$ by Student's $t$ test unless otherwise specified.

A

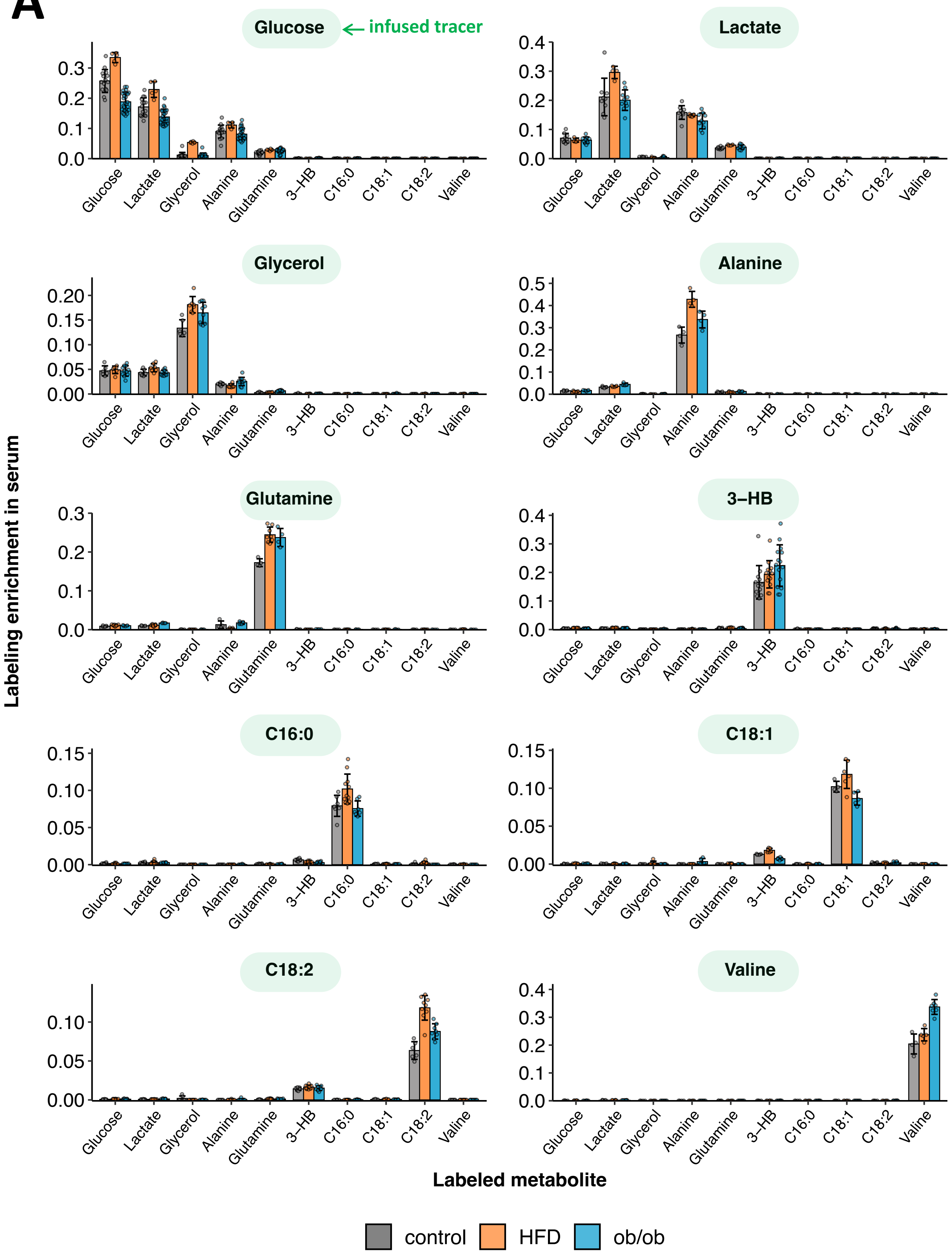


Figure S6 (continued)

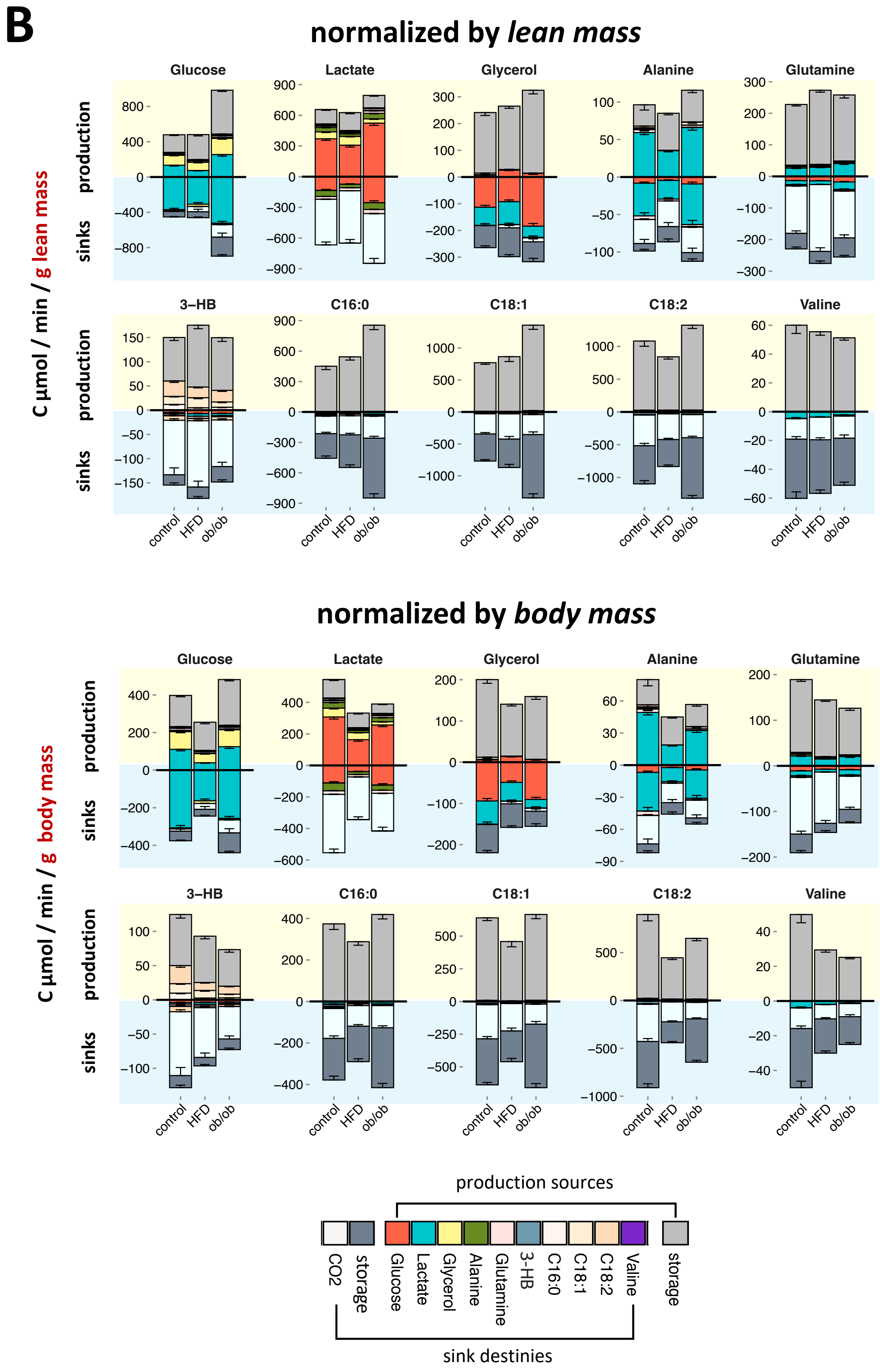


Figure S6 (continued)

**Figure S6. Flux Analysis in Lean and Obese Mice. Related to Figure 5.**

**(A)** $^{13}$C-labeling (without normalization) of serum nutrients during the isotopic steady state of $^{13}$C nutrient infusions in control and obese mice. Data is presented as mean ± SD.

**(B)** Production and sink fluxes of circulating nutrients in the control and obese mice. Fluxes are normalized by lean mass (top) or body mass (bottom), respectively. Fluxes on a per animal basis refer to Figure 5B. Data is presented as mean ± SEM.

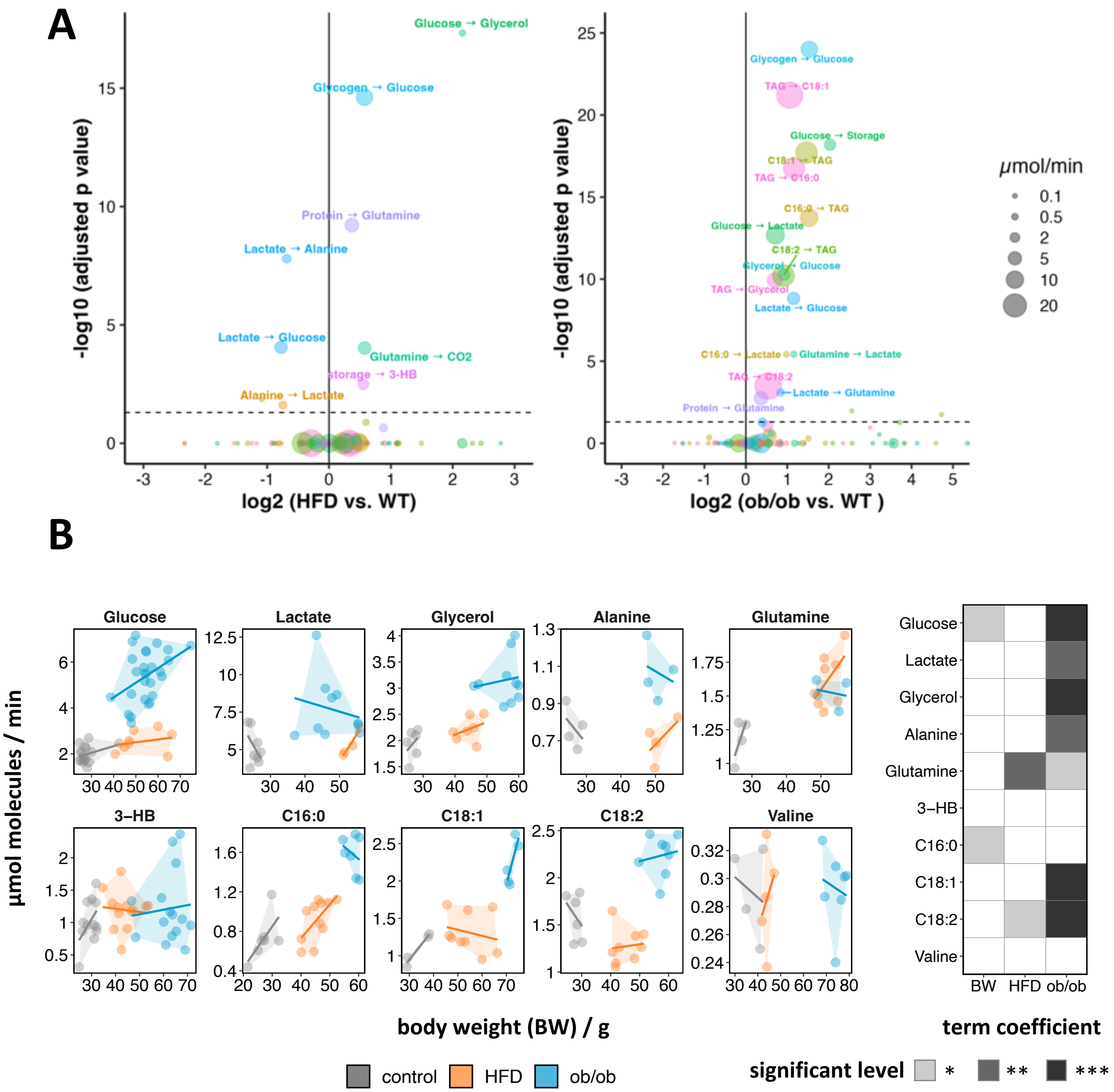


**Figure S7. Significance Analysis of Metabolic Fluxes Differences between the Control and Obese Mice. Related to Figure 5.**

**(A)** Volcano plot showing the fold change and associated *p* value of metabolic fluxes (per animal) in the obese mice vs. the control mice. The *p* values are adjusted using the Bonferroni method. The fluxes included in the plot correspond to those displayed in Figure 5B.

**(B)** Effect of body weight (BW) on the endogenous circulatory turnover fluxes $R_a$. The $R_a$ is calculated as $R_a = \frac{I(1-L)}{L}$, where $I$ is the infusion rate, and $L$ the labeling of the fully labeled isotopologue (Bartman et al., 2021; Hui et al., 2020). Statistical procedure follows that of CalR (Mina et al., 2018). Analysis of covariance (ANCOVA) is first performed, fitted with $F_{circ}=\beta_0 + \beta_1 BW + \beta_2 HFD' + \beta_3 ob/ob'$, where the BW is included as a covariant. The interaction terms

between BW and phenotypes are all insignificant, and thus not included in the model. In the ANCOVA analysis, if the BW coefficient $\beta_1$ is not significant, it is then excluded, and the model is fitted with analysis of variance (ANOVA) with $F_{circ}=\beta_0+\beta_1 HFD'+\beta_2 ob/ob'$. With respect to the dummy variables, for HFD group, HFD'=1, ob/ob'=0; for ob/ob group, HFD'=0, ob/ob' = 1; for the control mice, both terms are zero. The significance level of the coefficients is shown in the right-side heatmap. It shows that the increased fluxes in ob/ob mice are a result of intrinsic metabolic rewiring, instead of merely due to the larger body size. (Fluxes displayed in panel A and Figure 5B are calculated from compiled data and simulation, and cannot establish a flux-BW one-to-one relationship, and thus unable to be analyzed with ANCOVA.)
HFD mice were on high-fat diet for 3-4 months before experiment (most data), or for extended period (6-12 months) to gain a larger body mass (over 60 g).

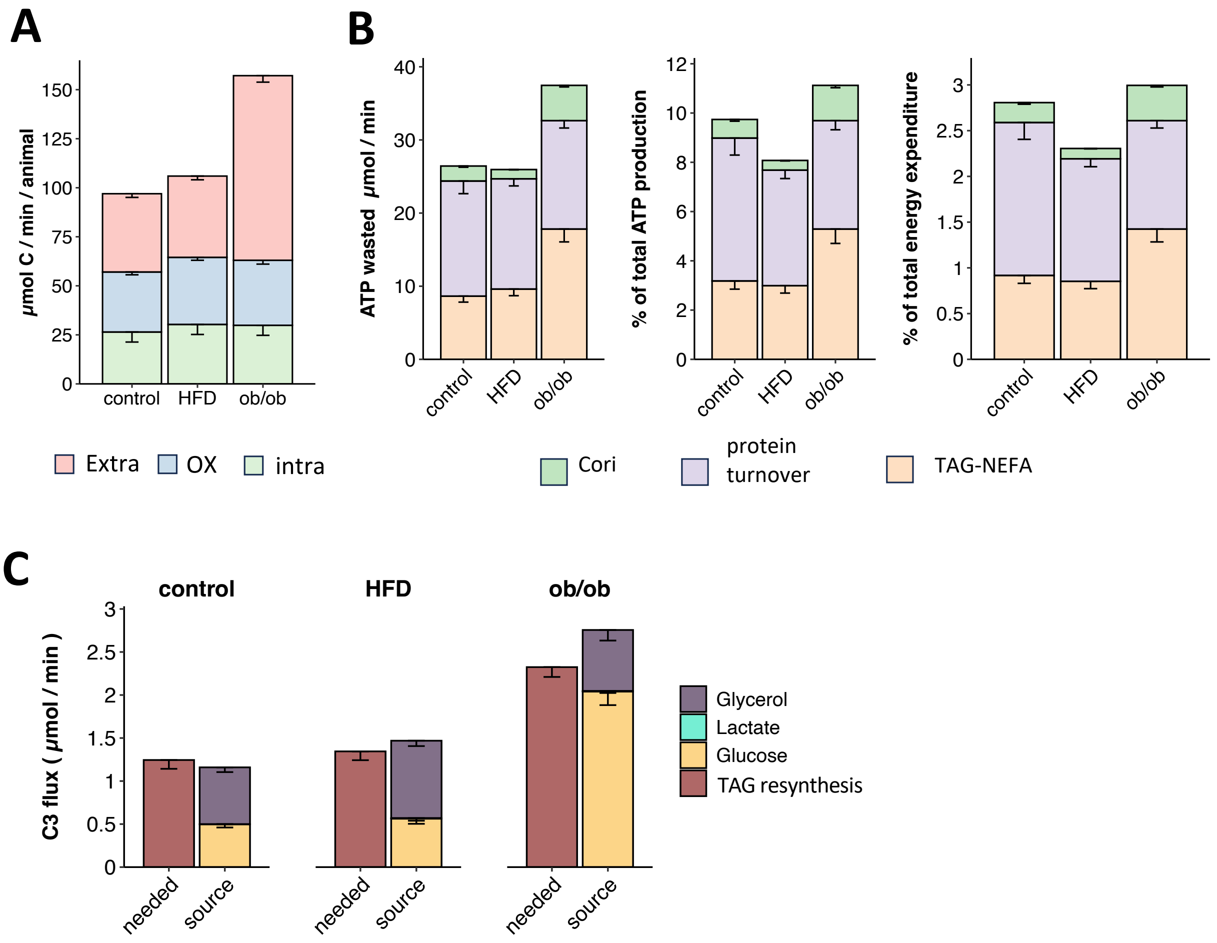


**Figure S8. Additional Fluxes Measured in the Obese Mice and Control Mice.**

**(A)** Fluxes of lipolysis-released fatty acids fated to extracellular reesterification (Extra), oxidation (OX), and intracellular reesterification (Intra).

**(B)** The amount of ATP wasted by the top 3 futile cycles.

**(C)** Comparison of the needed glycerol influx for triglyceride resynthesis with the storing fluxes of glycerol, lactate, and glucose.

All data is presented as mean ± SEM.
