## Supplemental Notes for "An Organism-Level Quantitative Flux Model of Energy Metabolism in Mice"

#### Supplementary Notes and Tables

Bo Yuan [1], Will Doxsey [1], Özlem Tok [1], Young Yon Kwon [1],

Karen E. Inouye [1,2], Gökhan S. Hotamışlıgil [1,2,3], Sheng Hui [1,2,4,*]

[1] Department of Molecular Metabolism, Harvard T. H. Chan School of Public Health, Boston, Massachusetts, USA.

[2] Sabri Ülker Center for Metabolic Research, Department of Molecular Metabolism, Harvard T.H. Chan School of Public Health, Boston, Massachusetts, USA.

[3] Broad Institute of Harvard and MIT, Cambridge, Massachusetts, USA.

[4] Lead Contact

**Supplementary Note 1.**

Calculation of Tracer-Derived 13CO2 Production Flux

**Supplementary Note 2.**

Calculation of Fluxes to Oxidation and Storages from a Circulating Nutrient

**Supplementary Note 3.**

Calculation of the Direct Fluxes to Oxidation and Storages from Circulating Nutrients

**Supplementary Note 4.**

Calculation of the Storage-Releasing Fluxes and Interconverting Fluxes of Circulating Nutrients

**Supplementary Note 5.**

Calculation of Fluxes of Circulating Nutrients from Storages as the Ultimate Source

**Supplementary Note 6.**

Calculation of Fluxes of Total Lipolysis and Intracellular Reesterification

**Supplementary Note 7.**

Estimating ATP Consumption in the Major Futile Cycles

**Supplementary Tables**

### Supplementary Note 1

#### Calculation of Tracer-Derived $^{13}CO_2$ Production Flux

| Notation | Definition | Unit |
|---|---|---|
| superscript $c$ | control mouse | |
| Superscript $t$ | $^{13}$C-tracer administered mouse | |
| Subscript $in$ | influx air into the metabolic cage | |
| Subscript $out$ | Outflux air out of the metabolic cage | |
| WVP | water vapor pressure | kPa |
| BP | barometric atmospheric pressure | kPa |
| $E$ | $^{13}$C enrichment (fraction) in the air $CO_2$ ($E_{in}, E_{out}$), or natural abundance in animal's endogenous nutrients ($E_{NA}$) | - |
| $C$ | $CO_2$ concentration in the air | ppm |
| $Q$ | Air flow through the metabolic cage | mL/min |
| $P$ | Animal's total ($^{12}$C, $^{13}$C) $CO_2$ production | mL/min |
| $J$ | $^{13}CO_2$ production by oxidation of the administered $^{13}$C-tracer | mL/min |
| $V$ | $^{12}CO_2$ production from by oxidation of animal's endogenous nutrient | mL/min |

Here we illustrate the procedure for calculating the production flux of $^{13}CO_2$ from a $^{13}$C-tracer. Such a procedure is necessary because the output of the stable isotope gas analyzer, $\delta^{13}$C, is not a flux quantity but one describing the amount of $^{13}$C atoms relative to the $^{12}$C atoms in the gas measured by the device. Moreover, this quantity reflects not only labeled $CO_2$ from the $^{13}$C-tracer administered to a mouse but also importantly labeled $CO_2$ from two other sources: natural abundance of $^{13}CO_2$ in the atmospheric air, and natural abundance of $^{13}$C in the animal's nutrient fuels that are being oxidized into $CO_2$. In the following, we will first convert $\delta^{13}$C to labeled fraction of $CO_2$ (or enrichment of $^{13}CO_2$) and then derive the equation for calculating tracer-derived $^{13}CO_2$ production flux.

**Calculation of Enrichment of $^{13}CO_2$ from $\delta^{13}$C**

The $^{13}CO_2$ level for an air sample is exported from the $^{13}CO_2$-analyzer in the form of $\delta\,^{13}$C, which is defined as

$$\delta\ ^{13}C = \left( \frac{\left(\frac{^{13}C}{^{12}C}\right)_{sample}}{\left(\frac{^{13}C}{^{12}C}\right)_{VPDB}} - 1 \right) \cdot 1000 \tag{1.1}$$

where $\left(\frac{^{13}C}{^{12}C}\right)_{sample}$ is the ratio of $^{13}$C abundance to the $^{12}$C abundance in the air sample, and VPDB (Vienna Pee Dee Belemnite) is the international reference standard for carbon isotopes, with $\left(\frac{^{13}C}{^{12}C}\right)_{VPDB} = 0.01123720.$

We define the enrichment of $^{13}CO_2$ in the sample as

$$E = \left( \frac{^{13}C}{^{12}C + {}^{13}C} \right)_{sample} \quad (1.2)$$

which can also be referred to as the labeled fraction of $CO_2$, a quantity easier for calculations later. Using Eq. 1.1, the enrichment can be expressed as a function of the measured $\delta^{13}$C,

$$E = \frac{\left( \frac{\delta^{13}C}{1000} + 1 \right) \cdot \left( \frac{^{13}C}{^{12}C} \right)_{VPDB}}{\left( \frac{\delta^{13}C}{1000} + 1 \right) \cdot \left( \frac{^{13}C}{^{12}C} \right)_{VPDB} + 1} \quad (1.3)$$

**A Metabolic Cage System Coupled to a Stable Isotope Gas Analyzer**

We next model our experimental setup of a metabolic cage system coupled to a stable isotope gas analyzer to derive the equation for calculating the production flux of $^{13}CO_2$ that originates from the $^{13}$C-tracer administered to a mouse. As shown in **Illustration 1**, in our experimental setup, the air was drawn with a pump at a known flow rate ($Q$) into each cage of a metabolic cage system. The outflux air passes through a stable isotope gas analyzer, which measures $\delta^{13}$C (or equivalently $CO_2$ enrichment $E_{out}$) and total $CO_2$ concentration ($C_{out}$). A cage can house a control animal or a $^{13}$C-tracer administered animal (denoted with superscript $c$ and $t$ in the measured quantities, respectively). The $CO_2$ concentration and enrichment measurements for an empty cage were taken as the $CO_2$ concentration ($C_{in}$) and enrichment ($E_{in}$) for the influx air.

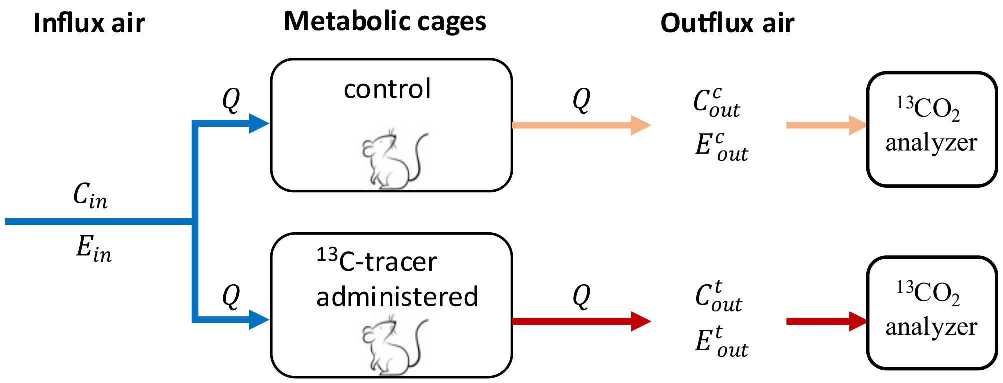


**Illustration 1.** Measurement of $^{13}CO_2$ using a metabolic cage system coupled to a stable isotope gas analyzer.

The directly measured $CO_2$ concentration $C'$ is corrected for water vapor pressure (WVP, in unit of kPa) in the air and non-standard barometric atmospheric pressure (BP, kPa). The corrected $CO_2$ concentration $C$ (ppm) in the air is calculated as

$$C = C' \cdot \frac{101.325}{BP - WVP}$$

##### $^{13}$C-Natural Abundance in the Animal's Endogenous Carbon Pool

To delineate the tracer-derived $^{13}CO_2$ production from natural abundance-derived $^{13}CO_2$ production, here we calculate the enrichment of exhaled $^{13}CO_2$ for a control mouse that does not receive $^{13}$C-tracer administration. For the mouse, the total $CO_2$ production flux $P^c$ (mL/min) can be expressed as

$$P^c = Q\ (C_{out}^c - C_{in})\ \cdot 10^{-6} \tag{1.4}$$

where $Q$ is the air flow rate at 2000 mL/min through the cage, $C_{in}$ and $C_{out}^c$ are the $CO_2$ concentration (ppm) measured in the air inflow and outflow, respectively, and the superscript *c* indicates the control group. Production flux of $^{13}CO_2$ from the control mouse is

$$P^c E_{NA} = Q\ (C_{out}^c\ E_{out}^c - C_{in}\ E_{in})\ \cdot 10^{-6} \tag{1.5}$$

where $E_{NA}$ is the $^{13}$C natural abundance of the endogenous nutrients used by the mouse, and $E_{in}$ and $E_{out}$ are the $^{13}CO_2$ enrichment measured in the air inflow and outflow, respectively. Combining Eqs. (1.4) and (1.5) gives natural $^{13}$C-enrichment $E_{NA}$ in the animal's endogenous carbon pool and exhaled $CO_2$ as

$$E_{NA} = \frac{C_{out}^c\ E_{out}^c - C_{in}E_{in}}{C_{out}^c - C_{in}} \tag{1.6}$$

##### Production of $^{13}CO_2$ by Oxidation of Administered $^{13}$C-Tracer

Similar to Eq. (1.4), for a mouse administered with $^{13}$C-tracers, the total $CO_2$ production $P^t$ (mL/min) can be written as

$$P^t = Q\ (C_{out}^t - C_{in})\ \cdot 10^{-6} \tag{1.7}$$

where the superscript $t$ indicates tracer-administered mouse. The total $CO_2$ production flux $P^t$ comprises the oxidation flux $V$ by burning the endogenous nutrients in the body, plus the oxidation flux $J$ by burning the administered tracer, and this gives

$$P^t = V + J \tag{1.8}$$

Similar to Eq. (1.5), a mass balances on the $^{13}CO_2$ for the $^{13}$C-tracer administered mouse gives

$$V \cdot E_{NA} + J = Q \cdot (C_{out}^t\ E_{out}^t - C_{in}\ E_{in})\ \cdot 10^{-6} \tag{1.9}$$

Integrating Eqs. (1.7-1.9) gives the tracer-derived $^{13}CO_2$ production rate as

$$J\ (mL/min) = \frac{Q}{1 - E_{NA}} \cdot [\ C_{out}^{t}\ (E_{out}^{t} - E_{NA}) - C_{in}\ (E_{in} - E_{NA})] \cdot 10^{-6} \qquad (1.10)$$

Converting the unit of $J$ from mL/min to μmol/min, we get

$$J\ (\mu mol/min) = \frac{Q}{22.4(1 - E_{NA})} \cdot [C_{out}^{t}\ (E_{out}^{t} - E_{NA}) - C_{in}\ (E_{in} - E_{NA})] \cdot 10^{-3} \qquad (1.11)$$

### Supplementary Note 2

#### Calculation of Fluxes to Oxidation and Storages from a Circulating Nutrient

| Notation | Definition | Unit |
|---|---|---|
| $f_{ox}$ | Fraction of oxidation | - |
| $J$ | $^{13}CO_2$ production by oxidation of the administered $^{13}$C-tracer | nmol/min/g |
| $I$ | $^{13}$C-tracer infusion rate | nmol C/min/g |
| $P^{total}$ | influx of total ($^{12}$C, $^{13}$C) glucose carbon into circulation | nmol C/min/g |
| $P$ | influx of endogenous ($^{12}$C) glucose carbon into circulation | nmol C/min/g |
| $P^*$ | influx of $^{13}$C recycle from labeled gluconeogenic substrates to glucose | nmol C/min/g |
| $R_a$ ($F_{circ}^{atom}$) | endogenous ($^{12}$C) rate of appearance (circulatory turnover) | nmol C/min/g |
| $J_{OX}$ | $CO_2$ production from oxidation of endogenous ($^{12}$C) glucose | nmol C/min/g |
| $J_S$ | Storage flux of endogenous ($^{12}$C) glucose | nmol C/min/g |
| $\phi$ | fraction of glucose $^{13}$C influx into circulation that is oxidized into $^{13}CO_2$ | - |
| $L$ | $^{13}$C-labeling in glucose | - |

Here in this note we derive the equation for calculating the flux from a circulating nutrient to its final oxidation product $CO_2$ ( $J_{OX}$) and nutrient storages in tissues ($J_S$). For simplicity, we use the circulating glucose as an example. The derivation however generally applies to any circulating nutrient.

At the steady state of $^{13}$C-glucose infusion, we have the fraction of oxidation $f_{ox}$ defined in the main text as the ratio of $^{13}CO_2$ production flux $J$ and the $^{13}$C-glucose infusion rate $I$, or

$$f_{ox} = \frac{J}{I} \tag{2.1}$$

To calculate the oxidation flux ( $J_{OX}$ ) originating from endogenous glucose, we introduce a quantity $\phi$ that describes the fraction of glucose carbon atoms (regardless of $^{12}$C or $^{13}$C) that are oxidized relative to the total $^{13}$C influx into circulation (infused + metabolically recycled; see Eq. 2.9), with

$$\phi = \frac{J}{P^{total}L} = \frac{If_{ox}}{P^{total}L} \tag{2.2}$$

$P^{total}$ is the total carbon influx (including both $^{12}$C and $^{13}$C) of glucose into circulation, and $L$ is the fraction of labeled carbon atoms in circulating glucose. $P^{total}L$ represents the total $^{13}$C glucose influx. The relation between the total glucose production $P^{total}$ and the endogenous production ($P$) of unlabeled carbon atoms is

$$P^{total}(1 - L) = P \tag{2.3}$$

Integrating Eqs. (2.2) and (2.3) gives

$$\phi = \frac{If_{ox}(1 - L)}{PL} \tag{2.4}$$

We can then have the flux of $CO_2$ generated from the endogenous $^{12}$C glucose production

$$J_{OX} = P\phi = R_a f_{ox} \tag{2.5}$$

where $R_a$ is the carbon-atom circulatory turnover flux for glucose, defined as

$$R_a = \frac{I(1-L)}{L} \tag{2.6}$$

Here $R_a$ is the same as $F_{circ}^{atom}$ defined in earlier works (Bartman et al., 2021; Bornstein et al., 2023; Hui et al., 2017, 2020). In like manner, the endogenous $^{12}$C flux $J_S$ of a circulating nutrient deposited into storages can be derived as

$$J_S = R_a(1 - f_{ox}) \tag{2.7}$$

This concludes our derivations for the $J_{OX}$ and $J_S$. Note it is clear from Eqs. (2.5) and (2.7) that the two quantities add up to the total carbon-atom circulatory turnover flux,

$$J_{OX} + J_S = R_a \tag{2.8}$$

##### A Side Note on $R_a$ versus Production Flux $P$

For most nutrients, $R_a$ is numerically close to the calculated endogenous $^{12}$C production flux $P$ (**Figure 4B**; derived in **Suppl. Note 4**). In glucose and lactate, however, $R_a$ is about 15% smaller than $P$. This discrepancy arises from recycling of carbon atoms in these molecules. Below is a discussion of this difference, again using $^{13}$C-glucose infusion as an example.

As shown in Illustration 2, total glucose production $P^{total}$ and comprises three streams, and can be expressed as

$$P^{total} = P + P^* + I \tag{2.9}$$

where $P^*$ is the flux of recycling of $^{13}$C atoms from labeled gluconeogenic substrates back to the glucose pool. $P^* + I$ constitutes the total $^{13}$C influx into circulating glucose, and accounts for the labeled fraction $L$. Combining Eqs. (2.3) and (2.9), we get

$$\frac{P}{1-L} = P + P^* + I \tag{2.10}$$

Rearrangement of this equation gives

$$P = I\frac{1-L}{L} + P^*\frac{1-L}{L} = R_a + P^*\frac{1-L}{L} \tag{2.11}$$

For most nutrients, the isotope recycling $P^*$ (infused X → nutrient Y → … → circulating X) is negligible, and results in $P = R_a$. For infused $^{13}$C-glucose and lactate, and alanine to a less extent, however, the recycling $P^*$ (e.g., though the Cori cycle) is quantitatively significant, and therefore renders $P > R_a$.

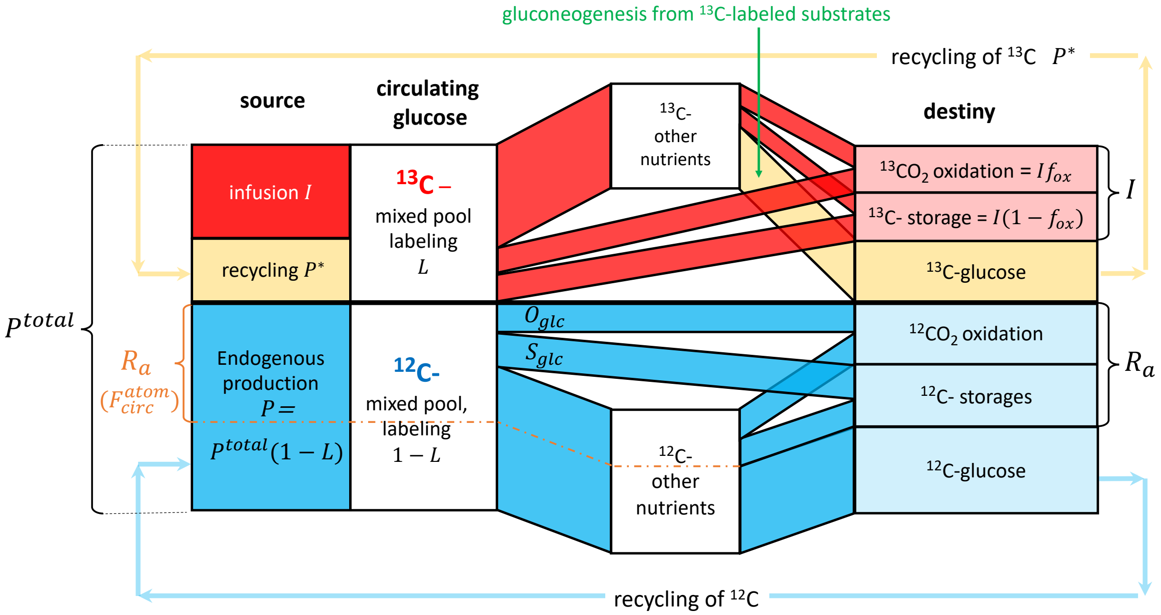


**Illustration 2.** The production and consumption of $^{12}$C and $^{13}$C of glucose during the steady state of $^{13}$C glucose infusion.

The above discussion shows a subtle but important difference between $R_a$ and $P$. Even though both terms are used to describes the production flux (release into circulation), $R_a$ does not count recycled atoms, and only count the "fresh new" ones as appearance. For example, if a glucose atom X entering into the blood stream is gluconeogenically from an atom in lactate, which can be traced back to the same atom in glucose in a previous metabolic cycle, then this atom X is not counted into $R_a$. $P$ on the other hand does count such recycled atoms, recognizing them as a *bona fide* component of the nutrient's influx into circulation.

With this understanding, it makes sense that the sum of a nutrient's destiny fluxes to oxidation and storages is equal to the "net" production flux $R_a$, as shown in Eq. (2.8): it does not count those recycled atoms, as these atoms have not been eventually fated yet.

Many literatures measured basal glucose *disposal* rate using radioactive $^{14}$C-glucose as a tracer, with the disposal rate calculated as $I\ (dpm/min/g)\ /\ L\ (dpm/nmol)$ , with $L$ being the glucose specific activity in serum (Del Prato et al., 1993). This approach is essentially the same as Eq. (2.6) (recognizing negligible mass of the radioactive infusate, $I \ll R_a$). It is important to note that such disposal refers to the removal of $R_a$: glucose's metabolism to lactate is counted as a disposal only if that lactate is not later converted back to glucose.

### Supplementary Note 3

#### Calculation of ***<u>Direct</u>*** Fluxes to Oxidation and Storages from Circulating Nutrients

| Notation | Definition | Unit |
|---|---|---|
| subscript $j$ | labeled nutrient $j$ | - |
| superscript $k$ | infused tracer $k$ | - |
| $L_j^k$ | Labeling of nutrient $j$ under the infusion of tracer $k$ | - |
| $\boldsymbol{L_1}$ | Matrix of labeling $L_j^k$ | |
| $f_{ox}$ | Fraction of oxidation | - |
| $I$ or $I_k$ | Infusion rate of $^{13}$C-tracer $k$ | nmol C/min/g |
| $\boldsymbol{I}$ | Matrix of infusion rate | - |
| $O_j^{total}$ | Direct and total ($^{12}$C, $^{13}$C) oxidation flux of nutrient $j$ | nmol C/min/g |
| $S_j^{total}$ | Direct and total ($^{12}$C, $^{13}$C) storage flux of nutrient $j$ | nmol C/min/g |
| $O_j$ | Direct oxidation flux of endogenous ($^{12}$C) nutrient $j$ | nmol C/min/g |
| $S_j$ | Direct storage flux of endogenous ($^{12}$C) nutrient $j$ | nmol C/min/g |
| $O_j^*$ | Direct oxidation flux of $^{13}$C in nutrient $j$ | nmol C/min/g |
| $S_j^*$ | Direct storage flux of $^{13}$C in nutrient $j$ | nmol C/min/g |
| $f_{C16:0}$ | Atom fraction of palmitate in NEFA | - |
| $X_{NEFA}$ ($S_{NEFA}$) | Extracellular reesterification (storage) flux ($^{12}$C) of NEFA | nmol C/min/g |
| $f_{Val}$ | Atom fraction of valine in protein | - |

In Note 2 we calculated the oxidation and storage fluxes from a circulating nutrient. These fluxes describe oxidation and storage as the ultimate but not direct destinies for the nutrient. This is because a nutrient can be converted into other circulating nutrients before being oxidized or stored. For example, circulating glucose can be converted into circulating lactate, which then feeds the tissue TCA cycle for oxidation. Thus, while most of the circulating glucose is eventually oxidized, almost negligible portion of it is directly oxidized. Here in this note, we aim to calculate the *direct* fluxes to oxidation and storage from circulating nutrients.

##### Calculation of the Direct Oxidation Flux

Again using glucose as an example, we consider the steady state of $^{13}CO_2$ production during a nonperturbative infusion of $^{13}$C-glucose, with an infusion rate $I$. For a circulating nutrient $j$, let $O_j^{total}$ be its direct oxidation flux (including both $^{12}$C and $^{13}$C), and $L_j$ its labeling in circulation. It follows that the direct $^{13}CO_2$ production flux $O_j^*$ from nutrient $j$ can be expressed as

$$O_j^* = O_j^{total} L_j \tag{3.1}$$

The $^{12}$C accounts for fraction $1 - L_j$ in nutrient $j$, and its direct oxidative flux $O_j$ is

$$O_j = O_j^{total}(1 - L_j) \tag{3.2}$$

As the $^{13}CO_2$ production flux by the mouse ($If_{ox}$) is equal to the sum of $O_j^*$ from all circulating nutrients that are labeled by the infused $^{13}$C-tracer (including circulating glucose), we get

$$\sum O_j^* = If_{ox} \tag{3.3}$$

Integrating the above three equations gives

$$\sum \frac{O_j}{(1 - L_j)} L_j = If_{ox} \tag{3.4}$$

While Eq. (3.4) is derived from the example of infusion of $^{13}$C-glucose, it applies to the infusion of other $^{13}$C-labeled nutrients. As such, the infusion of $N$ different $^{13}$C-nutrients gives a linear system of $N$ equations, or

$$\boldsymbol{A} \cdot \begin{bmatrix} O_1 \\ O_2 \\ \vdots \\ O_N \end{bmatrix} = \begin{bmatrix} I_1 f_{ox_1} \\ I_2 f_{ox_2} \\ \vdots \\ I_N f_{ox_N} \end{bmatrix} \tag{3.5}$$

$$with\ \boldsymbol{A} = \begin{bmatrix} \frac{L_1^1}{1 - L_1^1} & \frac{L_2^1}{1 - L_2^1} & \cdots & \frac{L_N^1}{1 - L_N^1} \\ \frac{L_1^2}{1 - L_1^2} & \frac{L_2^2}{1 - L_2^2} & \cdots & \frac{L_N^2}{1 - L_N^2} \\ \vdots & \vdots & \ddots & \vdots \\ \frac{L_1^N}{1 - L_1^N} & \frac{L_2^N}{1 - L_2^N} & \cdots & \frac{L_N^N}{1 - L_N^N} \end{bmatrix}$$

In matrix $\boldsymbol{A}$, each row corresponds to the infusion of a specific $^{13}$C-tracer, and each column corresponds to the labeling of a specific nutrient under infusion of different tracers. In other words, $L_j^k$ represents the labeling of the $j^{th}$ circulating nutrient under the infusion of the $k^{th}$ tracer.

Eq. (3.5) can be solved exactly for $O_j$. Due to measurement errors, however, the solutions can contain rare cases of negative values, albeit small ones. To avoid negative values, we use an optimization procedure that finds the set of values for $O_i$ that best match the measurements. Normally, the optimization procedure minimizes the sum of squares of residuals between the fitted values and observed values for the entries on the right side of Eq. (3.5). As the residuals are proportional to the infusion rate $I_k$ of tracer $k$, this procedure prioritizes optimization for nutrients with high infusion rates and generates poorly fitted results for nutrients with low infusion rates. To reduce the bias against small-flux nutrients, we normalized each row in matrix $\boldsymbol{A}$ respectively by $I_k$ , and renders the right-side into a vector of $f_{ox}$ values, or

$$(\boldsymbol{I}^{\#} \odot \boldsymbol{A}) \cdot \begin{bmatrix} O_1 \\ O_2 \\ \vdots \\ O_N \end{bmatrix} = \begin{bmatrix} f_{ox_1} \\ f_{ox_2} \\ \vdots \\ f_{ox_N} \end{bmatrix}, with\ \boldsymbol{I} = \begin{bmatrix} -\ \boldsymbol{i}_1^T\ - \\ -\ \boldsymbol{i}_2^T\ - \\ \vdots \\ -\ \boldsymbol{i}_N^T\ - \end{bmatrix} \tag{3.6}$$

where $-\ \boldsymbol{i}_k^T\ -$ is a transposed and horizontal vector of length-*N* of the same element $I_k$, # is defined as taking element-wise reciprocal in the matrix, and ⊙ taking element-wise product between two matrices. Solving Eq. (3.6) using the least square optimization method gives the endogenous oxidation fluxes of the circulating nutrients.

In addition, to ease calculation in the R script, the matrix $\boldsymbol{A}$ can be rewritten as

$$\boldsymbol{A} = \{\ \boldsymbol{L_1} \odot (\boldsymbol{1} - \boldsymbol{L_1})^{\#}\ \}\ \ with\ \boldsymbol{L_1} = \begin{bmatrix} L_1^1 & L_2^1 & \dots & L_N^1 \\ L_1^2 & L_2^2 & \dots & L_N^2 \\ \vdots & \vdots & \ddots & \vdots \\ L_1^N & L_2^N & \dots & L_N^N \end{bmatrix} \tag{3.7}$$

where **1** is a $N \times N$ matrix of 1's.

##### Calculation of the Direct Storing Flux

The direct storing fluxes of circulating nutrients can be derived in like manner as the direct oxidation flux mentioned above. For a circulating nutrient $i$ that is labeled by the infused tracer, this nutrient's direct $^{13}C$-storing flux $S_j^*$ can be expressed as

$$S_j^* = S_j^{total} L_j \tag{3.8}$$

where $S_j^{total}$ is the total direct storing flux (including both $^{12}C$ and $^{13}C$). As the $^{12}C$ accounts for fraction $1 - L_j$ in nutrient $j$, its direct endogenous storing flux $S_j$ can be expressed as

$$S_j = S_j^{total}(1 - L_j) \tag{3.9}$$

As the storing flux $I(1 - f_{ox})$ of the infused tracer is equal to the sum of $S_j^*$ from each labeled nutrient $j$, we get

$$\sum S_j^* = I(1 - f_{ox}) \tag{3.10}$$

Integrating Eqs. (3.8 - 3.10) gives

$$\sum \frac{S_j}{(1 - L_j)} L_j = I(1 - f_{ox}) \tag{3.11}$$

Integrating Eq. (3.11) from a set of infusions of $N$ $^{13}C$-nutrients gives a linear system

$$\boldsymbol{A}\cdot\begin{bmatrix}S_1\\S_2\\\vdots\\S_N\end{bmatrix}=\begin{bmatrix}I_1(1-f_{ox_1})\\I_2(1-f_{ox_2})\\\vdots\\I_N(1-f_{ox_N})\end{bmatrix} \tag{3.12}$$

To reduce the ordinary least squares-bias against small-flux nutrients, similar to Eq. (3.6), the direct storing fluxes $S_j$ are solved from the "normalized" matrix operation below

$$(\boldsymbol{I}^{\#}\odot\boldsymbol{A})\cdot\begin{bmatrix}S_1\\S_2\\\vdots\\S_N\end{bmatrix}=\begin{bmatrix}1-f_{ox_1}\\1-f_{ox_2}\\\vdots\\1-f_{ox_N}\end{bmatrix} \tag{3.13}$$

Summation of Eqs. (3.3) and (3.10) gives

$$\sum(O_j^*+S_j^*)=I \tag{3.14}$$

This mirrors the basic assumption that at steady state, the infused $^{13}$C atoms are balanced with their ultimate disposal by both oxidative and storing fluxes via the labeled circulating nutrients, while the interconversions among nutrients are not the ultimate disposal (graphically depicted in **Illustration 2**). Similarly, summation of Eqs. (3.5) and (3.12) gives

$$\boldsymbol{A}\cdot\begin{bmatrix}O_1+S_1\\O_2+S_2\\\vdots\\O_N+S_N\end{bmatrix}=\begin{bmatrix}I_1\\I_2\\\vdots\\I_N\end{bmatrix} \tag{3.15}$$

It shows that given the same infusion parameters and labeling matrix $\boldsymbol{A}$, $O_j+S_j$ are fixed numbers, and are independent of the $f_{ox}$ values.

##### Calculation of Direct Oxidation and Storing (Extracellular Re-esterification) Fluxes of NEFA

The direct oxidation flux of the total circulating non-esterified fatty acids (NEFA) $O_{NEFA}$ is calculated based on the direct oxidation fluxes of palmitate $O_{C16:0}$, oleate $O_{C18:1}$, and linoleate $O_{C18:2}$, and the atom fractions of the three fatty acids in total NEFA, $f_{C16:0}$, $f_{C18:1}$, and $f_{C18:2}$, respectively. The calculation proceeds as

$$O_{NEFA}=\frac{O_{C16:0}+O_{C18:1}+O_{C18:2}}{f_{C16:0}+f_{C18:1}+f_{C18:2}} \tag{3.16}$$

The direct storing flux of NEFA, i.e., the extracellular reesterification flux $X_{NEFA}$, is calculated similarly as

$$X_{NEFA}=\frac{S_{C16:0}+S_{C18:1}+S_{C18:2}}{f_{C16:0}+f_{C18:1}+f_{C18:2}} \tag{3.17}$$

**Calculation of Direct Oxidation and Storing Fluxes of Total Circulating Amino Acids**

The direct oxidation flux of total circulating amino acids $O_{amino\_acids}$ can be approximated based on the direct oxidation flux of valine $O_{val}$, and the average valine atom fraction $f_{val}$ in proteins on a carbon atom basis. It is calculated as

$$O_{amino\_acids} = \frac{O_{Val}}{f_{Val}} \quad (3.18)$$

The direct storing fluxes of total circulating amino acids $S_{amino\_acids}$ is calculated in like manner

$$S_{amino\_acids} = \frac{S_{Val}}{f_{Val}} \quad (3.19)$$

### Supplementary Note 4

#### Calculation of Storage-Releasing Fluxes and Interconverting Fluxes of Circulating Nutrients

| Notation | Definition | Unit |
|---|---|---|
| subscript $j$ | labeled nutrient $j$ | - |
| superscript $k$ | infused tracer $k$ | - |
| $L, L_j^k$ | Labeling of nutrient $j$ under the infusion of tracer $k$ | - |
| $\boldsymbol{L_2}$ | Matrix of labeling | - |
| $f_{ox}$ | Fraction of oxidation | - |
| $I, I_k$ | Infusion rate of $^{13}$C-tracer $k$ | nmol C/min/g |
| $P^{total}$ | Total ($^{12}$C, $^{13}$C) glucose production flux during infusion | nmol C/min/g |
| $P$ | Endogenous ($^{12}$C) glucose production flux | nmol C/min/g |
| $F_j^t$ | Total ($^{12}$C, $^{13}$C) glucose production from nutrient $j$ | nmol C/min/g |
| $F_j$ | Endogenous ($^{12}$C) glucose production from nutrient $j$ | nmol C/min/g |
| $F_S$ | Endogenous ($^{12}$C) glucose production from storage (e.g., glycogen) | nmol C/min/g |

In this note we show how we calculate for each circulating nutrient its contributing fluxes from its tissue storages and other circulating nutrients. Doing the same calculation for all the circulating nutrients considered will yield the interconverting fluxes between them.

**Releasing Flux from Glycogen and Conversion Fluxes from Circulating Nutrients to Glucose**

Using glucose as an example, we aim to calculate the releasing flux $F_S$ from the glucose storage glycogen to glucose, and the conversion fluxes from other circulating nutrients $F_j$ to glucose, such as lactate, glycerol, and amino acids. For this, consider the steady state during a nonperturbative $^{13}$C-glucose infusion (with an infusion rate of $I$). Under the infusion, the total glucose production flux $P^{total}$ (including both $^{12}$C and $^{13}$C) is equal to the sum of all incoming fluxes to glucose from glycogen and $n$ other circulating nutrients,

$$P^{total} = I + \sum_{j}^{n} F_j^t + F_S \qquad (4.1)$$

where $F_j^t = \frac{F_j}{1-L_j}$ with $L_j$ the labeling of nutrient $j$. Note that $F_j^t$ is the total flux from nutrient $j$ to glucose including both endogenous unlabeled flux $F_j$ and labeled flux $F_j^t L_j$. Now assuming negligible labeling in the glycogen storage given its significant pool size relative to the limited $^{13}$C-infusion time and rate, the mass balance of the labeled glucose pool (with the labeling denoted as $L_{glc}$) requires

$$\sum_{j}^{n} F_j^t L_j + I = P^{total} L_{glc} \qquad (4.2)$$

where the left-hand-side is the incoming $^{13}$C flux to glucose from different sources. Eq. (4.1) and the expression for $F_j^t$ can be plugged into Eq. (4.2) to form an equation with $F_j$'s and $F_S$ as the only unknowns,

$$I + \sum_j^n \frac{F_j}{1-L_j} L_j = \left( I + \sum_j^n \frac{F_j}{1-L_j} + F_s \right) L_{glc} \tag{4.3}$$

Re-arranging Eq. (4.3) gives

$$\sum_j^n \frac{L_{glc} - L_j}{1-L_j} F_j + F_s L_{glc} = I\left(1 - L_{glc}\right) \tag{4.4}$$

To reorganize the equation, for the second term regarding glycogen storage-releasing flux, $F_s L_{glc}$ can be replaced with $\frac{F_s}{1-L_s}(L_{glc} - L_s)$, with $L_s = 0$. The storage-releasing flux in its new format readily fits the summation $\sum$ operator as the $n+1$ term. Conceptually, the glycogen storage can be regarded as the $n+1$ nutrient. Therefore, the Eq. (4.4) can be rewritten as

$$\sum_j^{\mathbf{n+1}} \frac{L_{glc} - L_j}{1-L_j} F_j = I\left(1 - L_{glc}\right) \tag{4.5}$$

While Eq. (4.5) is derived using the infusion of $^{13}$C-glucose as an example, it holds true for infusion of any other $^{13}$C-nutrient $k$, except that $I = 0$, regarding the mass balance of circulating glucose. The generalized equation is

$$\sum_j^{n+1} \frac{L_{glc}^k - L_j^k}{1-L_j^k} F_j = X\left(1 - L_{glc}^k\right), \qquad X = \begin{cases} I, & tracer\ k = {}^{13}C\text{–}\ glucose \\ 0, & tracer\ k \neq {}^{13}C\text{–}\ glucose \end{cases} \tag{4.6}$$

where $L_j^k$ and $L_{glc}^k$ is the labeling of circulating nutrient $j$ and glucose under the infusion of $^{13}$C-tracer $k$, respectively. Note that the labeling is a variable dependent on the infused $^{13}$C-nutrient $k$, and we denote the labeling values with superscript $k$. In contrast, the endogenous $^{12}$C-conversion flux $F_j$ from $n$ circulating nutrients and storage-releasing flux $F_{n+1}$ to glucose are regarded as constants independent of tracers. Our assumption here is that the $^{13}$C-infusate is in small quantity, and does not induce negative feedback on the endogenous production; instead, the consumption accordingly elevates to accommodate the infused quantity.

With infusion of $^{13}$C glucose and $n$ other $^{13}$C-nutrients, $n+1$ equations in the format of Eq (4.6) can be constructed. Integrated together, this gives a linear system $\boldsymbol{B} \cdot \boldsymbol{f} = \boldsymbol{i}$, or

$$\boldsymbol{B} \cdot \begin{bmatrix} F_1 \\ F_2 \\ \vdots \\ F_n \\ F_s \end{bmatrix} = \begin{bmatrix} I(1 - L_{glc}) \\ 0 \\ \vdots \\ 0 \\ 0 \end{bmatrix} \tag{4.7}$$

with

$$\boldsymbol{B} = \begin{bmatrix} \frac{L_{glc}^{glc} - L_1^{glc}}{1 - L_1^{glc}} & \frac{L_{glc}^{glc} - L_2^{glc}}{1 - L_2^{glc}} & \cdots & \frac{L_{glc}^{glc} - L_n^{glc}}{1 - L_n^{glc}} & \frac{L_{glc}^{glc} - L_s^{glc}}{1 - L_s^{glc}} \\ \frac{L_{glc}^{1} - L_1^{1}}{1 - L_1^{1}} & \frac{L_{glc}^{1} - L_2^{1}}{1 - L_2^{1}} & \cdots & \frac{L_{glc}^{1} - L_n^{1}}{1 - L_n^{1}} & \frac{L_{glc}^{1} - L_s^{1}}{1 - L_s^{1}} \\ \vdots & \vdots & \ddots & \vdots & \vdots \\ \frac{L_{glc}^{n} - L_1^{n}}{1 - L_1^{n}} & \frac{L_{glc}^{n} - L_2^{n}}{1 - L_2^{n}} & \cdots & \frac{L_{glc}^{n} - L_n^{n}}{1 - L_n^{n}} & \frac{L_{glc}^{n} - L_s^{n}}{1 - L_s^{n}} \end{bmatrix}$$

The 1st row corresponds to the $^{13}$C-glucose infusion, and the 2nd to the $(n+1)^{th}$ row corresponds to the infusion of $n$ other different $^{13}$C-nutrients, respectively. The 1st to the $n^{th}$ column contains the labeling of the $n$ circulating nutrients, respectively, and the $(n+1)^{th}$ column contains the labeling of glycogen storage, with $L_s^k = 0$ . Both $\boldsymbol{B}$ and $\boldsymbol{i}$ are known quantities from measurements, and $\boldsymbol{f}$ can be solved, subjected to $\boldsymbol{f} \geq \boldsymbol{0}$.

To simplify the calculations, the matrix $\boldsymbol{B}$ can be rewritten as

$$\boldsymbol{B} = \left( \begin{bmatrix} - \boldsymbol{l}^{T}{}_{glc}^{glc} - \\ - \boldsymbol{l}'^{T}{}_{glc}^{1} - \\ \vdots \\ - \boldsymbol{l}^{T}{}_{glc}^{n} - \end{bmatrix} - \boldsymbol{L_2} \right) \odot (\boldsymbol{1} - \boldsymbol{L_2})^{\#} \quad with\, \boldsymbol{L_2} = \begin{bmatrix} L_1^{glc} & L_2^{glc} & \cdots & L_n^{glc} & 0 \\ L_1^1 & L_2^1 & \cdots & L_n^1 & 0 \\ \vdots & \vdots & \ddots & \vdots & \vdots \\ L_1^n & L_2^n & \cdots & L_n^n & 0 \end{bmatrix} \qquad (4.8)$$

$- \boldsymbol{l}^{T}{}_{glc}^{k} -$ is the vector transposed, comprising the same element $L_{glc}^k$ with length of $n$+1, and **1** is the matrix of 1's with dimension ($n$+1) × ($n$+1). Here the first $n$ columns of $\boldsymbol{L_2}$ (excluding the last $\boldsymbol{0}$ column) is a subset of matrix $\boldsymbol{L_1}$ defined in Eq. (3.7). Summation of the $\boldsymbol{f}$ elements gives the total endogenous glucose production flux $P_{glu}$, with

$$P_{glc} = \sum_{j}^{n+1} F_j = \sum_{j}^{n} F_j + F_s \qquad (4.9)$$

**Generalization to Calculate the Production Flux of Any Circulating Nutrient**

The above calculation for the production fluxes of glucose can be generalized for any other circulating nutrient. For a target nutrient (TN) of interest, Eq. (4.6) becomes

$$\sum\nolimits_{j}^{n+1} \frac{L_{TN}^{k} - L_j^{k}}{1 - L_j^{k}} F_j = X\left(1 - L_{TN}^{k}\right), \qquad X = \begin{cases} I, & tracer\ k = {}^{13}C\text{–}TN \\ 0, & tracer\ k \neq {}^{13}C\text{–}TN \end{cases} \qquad (4.10)$$

where $L_j^k$ and $L_{TN}^k$ is the labeling of circulating nutrient $j$ and TN under the infusion of $^{13}$C-tracer $k$, respectively. Note that now $F_j$ is the production flux from nutrient $j$ or storage to TN. Eq. (4.8) can be rewritten in the form of a matrix operation, similar to Eq. (4.7), which can be solved for $F_j$. Looping through the matrix computation $n$ times for each of the other $n$ circulating nutrients as the TN, respectively, generates the total production and the interconversion fluxes among the circulating nutrients.

### Supplementary Note 5

#### Calculation of Fluxes of Circulating Nutrients from Storages as the ***Ultimate*** Source

| Notation | Definition | Unit |
|---|---|---|
| $P_{nutrient}$ | Endogenous ($^{12}$C) production flux of a circulating nutrient | nmol/min/g |
| $F_{A \to B}$ | Endogenous ($^{12}$C) flux from source *A* to produce *B* | nmol/min/g |
| subscripts | *S*, storage; *G*, glycogen; *T*, triglyceride; *P*, protein | - |
| $f_{G\,j}$ | Ultimate contribution *fraction* from glycogen to produce nutrient $j$ | - |
| $U_{G_j}$ | Ultimate contribution *flux* ($^{12}$C) from glycogen to produce nutrient $j$ | nmol/min/g |
| TN | target nutrient | - |

During fasting state, the storage pools are the *ultimate* source of circulating nutrients. For instance, circulating glucose can be derived from glycogen and circulating glycerol, while the latter is derived from triglyceride. As such, glycogen and triglyceride, but not glycerol, are the ultimate sources of circulating glucose. In this note, we aim to calculate such ultimate contribution fluxes from glycogen, triglyceride, and protein, to circulating nutrients.

To illustrate the calculation, here we calculate the ultimate contribution from glycogen. Take the production of glucose $P_{glc}$ as an example, which comprises the releasing flux from its storage glycogen $F_{s \to glc}$, and the conversion fluxes $F_{j \to glc}$ from circulating nutrients, or

$$F_{s \to glc} + \sum F_{j \to glc} = P_{glc} \tag{5.1}$$

Let $f_{G\,j}$ be the ultimate contribution fraction from glycogen to circulating nutrient $j$. Then $F_{j \to glc} f_{G\,j}$ is the flux of glycogen-derived carbons from nutrient $j$ to glucose. Summation of such direct fluxes from all contributing sources to circulating glucose, or $F_{s \to glc} + \sum F_{j \to glc}\, f_{G\,j}$, gives the total flow of glycogen carbons to circulating glucose. At steady state, this quantity is equal to the outgoing flux of the glycogen-derived glucose carbons, $P_{glc} f_{G\,glc}$, where $f_{G\,glc}$ is the glycogen's ultimate contribution fraction to glucose. Mathematically,

$$F_{s \to glc} + \sum F_{j \to glc}\, f_{G\,j} = P_{glc} f_{G\,glc} \tag{5.2}$$

Eq. (5.2) can be conceptually regarded as a mass balance equation for the labeled glucose using $^{13}$C-glycogen as the tracer, with $f_{G_j}$ as the labeled fraction of nutrient $j$.

Similarly, for circulating alanine, as another example, its production flux $P_{ala}$ comprises the releasing flux from storage, the proteins, and the conversion from other circulating nutrients

$$F_{s\to ala} + \sum F_{j\to ala} = P_{ala} \tag{5.3}$$

regarding glycogen-derived carbon atoms, we get

$$F_{s\to ala} \cdot 0 + \sum F_{j\to ala} f_{G_i} = P_{ala} f_{G_{ala}} \tag{5.4}$$

To generalize Eqs. (5.2) and (5.4) to any target circulating nutrient (TN), with respect to glycogen's contribution, we have

$$F_{s\to TN}\, a_{TN} + \sum F_{j\to TN}\, f_{G\,j} = P_{TN}\, f_{G\,TN} \tag{5.5}$$

$$\text{with } a_{TN} = \begin{cases} 1, & TN = glucose\ or\ lactate \\ 0, & TN \neq glucose\ or\ lactate \end{cases}$$

Eq. (5.5) can be rearranged with terms containing the unknown $f_G$ on the left side of the equation, and known quantities on the right side, such that

$$-P_{TN}\, f_{G\,TN} + \sum F_{j\to TN}\, f_{G\,j} = -F_{s\to TN}\, a_{TN} \tag{5.6}$$

Integrating Eq. (5.6) across all circulating nutrients, coded with subscript 1, 2…to $n$, we get a linear system

$$\boldsymbol{M} \cdot \begin{bmatrix} f_{G\,1} \\ f_{G\,2} \\ \vdots \\ f_{G\,n} \end{bmatrix} = \begin{bmatrix} -F_{s\to 1} a_1 \\ -F_{s\to 2} a_2 \\ \vdots \\ -F_{s\to n} a_n \end{bmatrix} \tag{5.7}$$

where $\boldsymbol{M}$ is defined as

$$\boldsymbol{M} = \begin{bmatrix} -P_1 & F_{2\to 1} & \cdots & F_{n\to 1} \\ F_{1\to 2} & -P_2 & \cdots & F_{n\to 2} \\ \vdots & \vdots & \ddots & \vdots \\ F_{1\to n} & F_{2\to n} & \cdots & -P_n \end{bmatrix}$$

All the inter-converting fluxes (e.g., $F_{1\to 2}$), storage releasing fluxes (e.g., $F_{s\to 1}$), and the total production fluxes (e.g., $P_1$) are known from previous calculations in Note 4. Thus, (5.7) can be solved for the ultimate contribution fractions from glycogen $f_{G\,j}$.

The ultimate contribution flux $U_G$ from glycogen to circulating nutrient $j$ is then calculated as

$$U_{G\,j} = P_j\, f_{G\,j} \tag{5.8}$$

Similarly, the ultimate contribution fraction of triglyceride $f_T$ to $n$ circulating nutrients can be acquired by solving

$$
\boldsymbol{M} \cdot \begin{bmatrix} f_{T_1} \\ f_{T_2} \\ \vdots \\ f_{T_n} \end{bmatrix} = \begin{bmatrix} -F_{s\to 1}\, b_1 \\ -F_{s\to 2}\, b_2 \\ \vdots \\ -F_{s\to n}\, b_n \end{bmatrix} \tag{5.9}
$$

$$
b_j = \begin{cases} 1, & j = C16{:}\,0, C18{:}\,1, C18{:}\,2, glycerol, or\ 3-HB \\ 0, & j \neq C16{:}\,0, C18{:}\,1, C18{:}\,2, glycerol, or\ 3-HB \end{cases}
$$

And the contribution fraction of proteins $f_P$ can be solved by

$$
\boldsymbol{M} \cdot \begin{bmatrix} f_{P_1} \\ f_{P_2} \\ \vdots \\ f_{P_n} \end{bmatrix} = \begin{bmatrix} -F_{s\to 1}\, c_1 \\ -F_{s\to 2}\, c_2 \\ \vdots \\ -F_{s\to n}\, c_n \end{bmatrix}, \qquad c_j = \begin{cases} 1, & j = amino\ acids \\ 0, & j \neq amino\ acids \end{cases} \tag{5.10}
$$

### Supplementary Note 6

#### Calculation of Fluxes of Total Lipolysis and Intracellular Reesterification Fluxes of Fatty Acids

The total lipolysis flux $P_{lipolysis}$ is calculated as 3-fold of glycerol production flux $P_{glycerol}$, with

$$
P_{lipolysis} = 3 \cdot P_{glycerol} \tag{6.1}
$$

The discrepancy between $P_{lipolysis}$ and the production flux of NEFA $P_{NEFA}$ unveils the intracellular re-esterification flux $I_{NEFA}$, i.e., the immediate fixation and recycling of lipolysis-produced fatty acids before they are release into circulation (Kalderon et al., 2000).

$$
I_{NEFA} = P_{lipolysis} - P_{NEFA} \tag{6.2}
$$

By Eq. (3.17) in *Supplementary Note 3*, we calculated the extracellular reesterification flux $X_{NEFA}$, and oxidation flux $O_{NEFA}$. As such, the fatty acids produced by lipolysis can be quantitatively partitioned into three distinct fates, with

$$
P_{lipolysis} = I_{NEFA} + X_{NEFA} + O_{NEFA} \tag{6.3}
$$

NEFA can be used for production of ketone bodies, but the related flux is very small compared with the NEFA production fluxes, and these fluxes are neglected when calculating the NEFA reesterification fluxes.

### Supplementary Note 7

#### Estimating ATP Consumption in the Major Futile Cycles

**ATP Counts in Cori cycle**

*a) From 1 glucose molecule to 2 pyruvate molecules:* glucose → glucose-6-phosphate, −1 ATP; fructose-6-phosphate → fructose-1,6-bisphosphate, −1 ATP; 1,3-bisphosphoglycerate → 3-phosphoglycerate, + 2ATP; phosphoenolpyruvate → pyruvate, + 2ATP. A total of 2 ATP molecules are produced from 1 glucose molecule.

*b) From 2 pyruvate molecules to 1 glucose molecule: pyruvate* → oxaloacetate, −2 ATP; oxaloacetate → phosphoenolpyruvate, − 2 GTP; 3-phosphoglycerate → 1,3-bisphosphoglycerate, − 2 ATP. A total of 6 ATP molecules are consumed to make one molecule of glucose.

c) Inter-conversion between pyruvate and lactate at equal flux involves no net energy change.

In total, **Cori cycle between glucose and lactate consumes a net of 4 ATP molecules per glucose molecule**.

**ATP Counts in TAG resynthesis**

For estimation purpose, we assume in this work that glucose is the major source of glycerol backbone in the TAG resynthesis (though our data suggests that glucose alone is not necessarily sufficient). In the calculation, we consider that TAG is resynthesized directly from NEFA and glucose-derived glycerol-3-phosphate (G3P) in adipose tissue.

a) Glycolysis in adipose tissue generate dihydroxyacetone 3-phosphate (DHAP), which is further reduced into G3P to support TAG synthesis. This procedure consumes 1 ATP (in the first two ATP-consuming steps in glycolysis) and one cytosolic NADH (counted as 2 ATP) per G3P molecule.
b) Three free fatty acids are activated into fatty acyl-CoA at the cost of 6 ATP. The G3P and acyl-CoA are then packed into TAG via the G3P pathway.

This way, the resynthesis of each TAG molecule via the glycolysis and G3P pathway consumes 9 ATP molecules.

**ATP Counts in Protein Turnover**

The synthesis of each peptide bond consumes 4 ATP molecules (Bender, 2012). The rate of ATP consumption in the protein turnover is calculated as

$$\frac{S_{val}}{5} f^{m}{}_{val} \cdot 4 \qquad (7.2)$$

where $S_{val}$ is the direct storing flux of valine back to protein, 5 is the number of carbon atoms in one valine molecule, and $f^m$ is the average fraction of valine in protein on a molecular basis.

##### Total Energy Expenditure (TEE)

TEE (calorie/min) is calculated using the Modified Weir equation (Mehta et al., 2015), based on the rate of $CO_2$ production $V_{CO_2}$, and $O_2$ consumption $V_{O_2}$ (L/min)

$$TEE = \left(3.941V_{O_2} + 1.106\,V_{CO_2}\right) \cdot 1000 \tag{7.3}$$

##### Rate of Total ATP Production $P_{ATP}$

The flux of total ATP production $P_{ATP}$ is calculated as

$$P_{ATP} = \sum O_i N_i \tag{7.4}$$

where $O_i$ is the direct oxidation carbon flux of circulating nutrients, and $N_i$ the number of ATP molecule produced by oxidation of per carbon atom of nutrient-$i$, taking the value of 5.3, 6.5, 5.5, and 5, for carbohydrates, fatty acids, ketone bodies, and amino acids, respectively (Bender, 2012).

The amount of energy released by ATP hydrolysis is estimated by the Gibbs free energy $\Delta G_{physiology}$ at a representative physiological condition: 37 °C, 5 mM ATP, 5 mM phosphate, and 0.5 mM ADP:

$$ATP + H_2O \rightleftharpoons ADP + P_i$$

$$\Delta G_{physiology} = \Delta G_{standard} + RTln\left(\frac{[ADP][P_i]}{[ATP]}\right) \tag{7.5}$$

where $\Delta G_{standard}$ = -31.5 KJ/mol (Meurer et al., 2017), the universal gas constant $R$ = 0.008314 KJ · (mol · K)$^{-1}$, $T$ = 310 K, and substrate concentration in the unit of moles. $\Delta G_{physiology}$ is calculated to be -51 KJ/mol or -12.2 kcal/mol (Beals et al., 1999).

The cellular concentration of ATP, ADP and Pi can vary across a wide range. Below we show the amount of energy (as absolute value) released by ATP hydrolysis under a range of substrate concentrations, with [ATP] and [Pi] varied from 1 to 10 mM, and [ADP] from 0.1 to 1 mM.

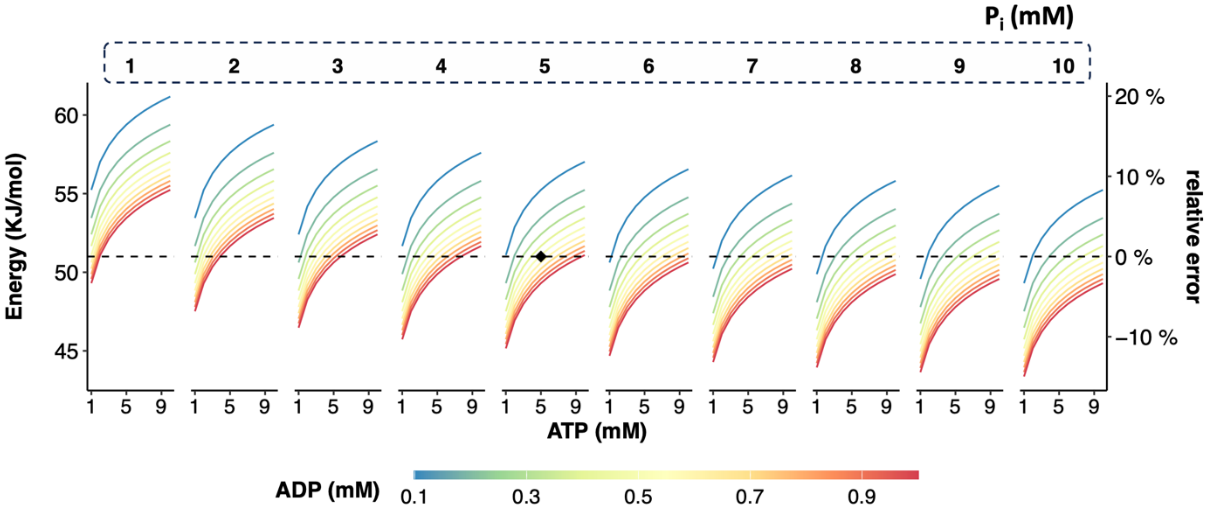


Despite 10-fold variance in concentration in the three substrates, the error is capped within a maximum of 20% compared with the reference value of 51 KJ/mol. While the Gibbs free energy also depend on intracellular pH and magnesium concentration, the associated variation is also within a reasonable narrow range (Rosing and Slater, 1972).

### Supplementary Tables

**Table S1. $^{13}C$-nutrient infusion parameters.**

| tracer | $^{13}CO_2$ measurement * | | | Serum labeling analysis | |
|---|---|---|---|---|---|
| | conc. mM | I.V. Bolus | Infusion | Infusion | |
| | | µL/g (*nmol/g*) | µL/g/min (*nmol/g/min*) | conc. mM | µL/g/min (*nmol/g/min*) |
| Glucose | 16.5 | 4 - 8 (*66 - 132*) | 0.04 - 0.08 (*0.66 - 1.32*) | 200 | 0.08 - 0.1 (*16 - 20*) |
| Lactate | 12.3 | 4 - 8 (*49.2 - 98.4*) | 0.04 - 0.08 (*0.492 - 0.984*) | 380 | 0.08 - 0.1 (*30.4 - 38*) |
| Glycerol | 13.8 | 4 - 8 (*55.2 - 110.4*) | 0.04 - 0.08 (*0.552 - 1.104*) | 110.1 | 0.1 (*11.01*) |
| Alanine | 8 | 4 - 8 (*32 - 64*) | 0.04 - 0.08 (*0.32 - 0.64*) | 103.7 | 0.1 (*10.37*) |
| Glutamine | 6.1 | 4 - 8 (*24.4 - 48.8*) | 0.04 - 0.08 (*0.244 - 0.488*) | 85.9 | 0.1 (*8.59*) |
| 3-HB | 12.5 | 4 - 8 (*50 - 100*) | 0.04 - 0.08 (*0.5 - 1*) | 61.5 | 0.08 - 0.1 (*4.92 - 6.15*) |
| Acetate | 14.4 | 4 - 8 (*57.6 - 115.2*) | 0.04 - 0.08 (*0.576 - 1.152*) | 100.8 | 0.1 (*10.08*) |
| C16:0 | 3.6 | 4 - 8 (*14.4 - 28.8*) | 0.04 - 0.08 (*0.144 - 0.288*) | 10.7 | 0.2 (*2.14*) |
| C18:1 | 4.8 | 4 - 8 (*19.2 - 38.4*) | 0.04 - 0.08 (*0.192 - 0.384*) | 10 | 0.28 - 0.35 (*2.8 - 3.5*) |
| C18:2 | 4.8 | 4 - 8 (*19.2 - 38.4*) | 0.04 - 0.08 (*0.192 - 0.384*) | 10 | 0.35 (*3.5*) |
| Valine | 18.4 | 4 - 8 (*73.6 - 147.2*) | 0.04 - 0.08 (*0.736 - 1.472*) | 20 | 0.1 (*2*) |
| Methionine | 16.2 | - | 0.08 (*1.296*) | - | |
| Tryptophan | 5.8 | - | 0.08 (*0.464*) | - | |

* For $^{13}CO_2$ measurement, the bolus dosage was generally 200 µL/animal, and the infusion rate was 2 µL/animal/min, for both lean and obese mice. Given the minimal dosage used, $^{13}CO_2$ production remains linear to a large range of administered dosage.

**Table S2. The ratio of $^{13}C$ labeling in the arterial blood and the tail-snip blood.**

Data is presented as mean ± SEM (n).

| infused tracer | phenotype | Glucose | Lactate | Alanine | Glutamine |
|---|---|---|---|---|---|
| Glucose | WT | 1.09 ± 0.03 (5) | 1.38 ± 0.03 (5) | 1.21 ± 0.07 (5) | 1.06 ± 0.03 (5) |
| Glucose | ob/ob | 1.02 ± 0.01 (7) | 1.09 ± 0.03 (7) | 1.11 ± 0.05 (5) | 1.12 ± 0.04 (7) |
| Lactate | WT | 1.07 ± 0.01 (6) | 1.94 ± 0.13 (6) | 1.42 ± 0.09 (6) | 1.13 ± 0.02 (6) |
| Lactate | ob/ob | 1.06 ± 0.02 (5) | 1.63 ± 0.12 (5) | 1.52 ± 0.17 (5) | 1.22 ± 0.04 (5) |
| Alanine | WT | 1 ± 0.05 (3) | 1.48 ± 0.14 (3) | 1.34 ± 0.1 (3) | 1.11 ± 0.04 (3) |
| Alanine | ob/ob | 1.06 ± 0.04 (3) | 1.31 ± 0.06 (3) | 1.4 ± 0.02 (3) | 1.2 ± 0.01 (2) |
| Glutamine | WT | 1.02 ± 0.07 (3) | 1.42 ± 0.05 (3) | - | 1.09 ± 0.1 (2) |
| Glutamine | ob/ob | 0.99 ± 0.01 (3) | 1.32 ± 0.08 (3) | 1.36 ± 0.07 (3) | 1.13 ± 0.03 (3) |

*note:* HFD-fed mice (but not the other 3 phenotypes) consistently had poor post-operational recovery from double catheter implantation. Thus, for HFD group, only jugular vein catheter was implanted, and tail-snip blood was taken for analysis. The arterial-venous ratio is assumed the same as WT mice.
